# Social wound care dynamics in an army ant society

**DOI:** 10.1101/2025.09.22.677721

**Authors:** Juan J. Lagos-Oviedo, Dharanish Rajendra, Chaitanya S. Gokhale, Thomas Schmitt, Erik T. Frank

**Author notes:** Corresponding authors: Juan J. Lagos-Oviedo (J.J.L.O.) and Erik T. Frank (E.T.F.), **Email**. **Competing Interest Statement:** We declare no competing interests.

## Abstract

Injuries and infections pose a significant threat to fitness. Animals cope with this problem by performing self-medication or receiving social care from members of the group. While the benefits of wound care in increasing survival are clear, the forces driving the evolution of these behaviours are not fully understood. Contrary to the current hypothesis that social wound care is more likely to evolve in small groups due to the high relative value of an individual, we demonstrate that group size is not necessary for its evolution or retention. Instead, our theoretical model formalises how injury rate and lethality are fundamental drivers of care rather than group size. This is further evidenced by the presence of wound care in the army ant *Eciton burchellii*, a species with colonies of ∼1 million ants that hunts pugnacious prey. Here, wound care significantly increased survival in ants with infected wounds, occurring in two phases: direct care at the raiding site, and treatment with antimicrobial secretions in the bivouac. This species is the first to provide on-site care, thereby minimising the latency to receive care and thus potentially the likelihood of lethal wound infection. Our findings reject the notion that small group size is necessary for the evolution or retention of social wound care, revealing instead that injury-driven care can be a widespread and adaptive feature of even the largest insect societies.

**Significance Statement:** Injuries and infections threaten survival, but animals can counter these dangers through self-medication or social care. It has been suggested that helping behaviours directed towards vulnerable or injured individuals, such as social wound care, are more likely to evolve in small groups. This study in an army ant with ∼1 million workers shows otherwise, with injury frequency likely being the key component for the evolution or retention of wound-care behaviours. The unique strategy of on-site care at the raiding site, followed by antimicrobial care inside the bivouac, significantly improved survival outcomes. Our theoretical model further shows that even at low rates of care, fitness benefits are high, suggesting that social wound care may be far more widespread than previously thought.

## Introduction

Animals are likely to acquire injuries during their life. These injuries are detrimental to individual fitness by affecting development (1, 2), foraging decision-making (3), locomotory performance (4–6), predation (7, 8) and infection risk (9, 10). Therefore, strategies to prevent and cope with injuries are expected to be under strong selection to reduce these costs. Animals prevent injuries by assessing risks during and before the hunt (4, 6), or during inter- and intraspecific competition (6, 11). However, when animals are injured, they can actively cope with the costs through two main strategies. First, they may engage in self-treatment by actively grooming at open wounds (wound care), applying antimicrobial compounds, or utilising antimicrobial plants (12–14).

Second, they can receive treatment by conspecifics, including antimicrobial wound care or even amputations to treat infections (10, 12–20). Social wound care behaviours have been documented across diverse taxa such as primates, ungulates, rodents, and ants (10, 12–24). For example, in the specialised termite-hunting species *Megaponera analis*, ants suffer injuries from fights with their prey (16). After a fight, injured workers emit a volatile help pheromone, eliciting nestmates to carry them back to the nest, where their wounds are subsequently treated (17, 25). This wound care behaviour reduces combat and infection mortality by 90% (17, 25). In carpenter ants, workers also receive wound care and nestmates even amputate injured legs inside the nest (15, 16). This prophylactic amputation behaviour reduces the mortality of experimentally infected workers by up to 88% after 72 hours (16).

Although there is growing evidence for the presence of social wound care behaviours in animals, we still lack a clear understanding of which ecological and evolutionary forces favour the evolution of this helping behaviour. The fact that social wound care has only been reported in species with relatively small group sizes (ranging from a handful in mammals to a few thousand individuals in ants) suggests that a high individual value, due to relatively low birth rates in these species, could favour the evolution of wound care (26). Additionally, the amount of non-lethal injuries incurred, such as from predator-prey interactions (1, 8, 27–29) or interspecific competition (11), may increase wound infection risks and therefore the necessity of caring for the injured. Thus, the relative individual value, the injury frequency and lethality may contribute to the evolution of social wound care behaviours, but to date, no study comprehensively examines their role in fitness.

To identify the importance of the individual value and injury rate on the evolution of social wound care, we studied this behaviour in the neotropical army ant *Eciton burchellii*. Similar to *M. analis*, in which social wound care has been reported (10, 17), the army ant *E. burchellii* raids pugnacious prey in large groups. But, in contrast to other wound-caring ant species (10, 16–18), army ants can reach colony sizes of over 1 million ants, which is one thousand times more than in *M. analis* (30–32). The relative value of individual workers to the colony is expected to decrease with increasing colony size, such that the loss of a single worker has a smaller impact on colony performance in larger societies. If this holds, then species with very large colonies, such as *E. burchellii*, should be less likely to exhibit social wound care behaviours. To formalise how interactions among group size, injury rate, lethality and social wound care translate into fitness, we also developed a general theoretical model applicable to *E. burchellii* and other group-living species. We further extend the model to examine how other aspects of social wound care that are not currently considered (25, 26), such as the latency to care and its efficacy, can buffer the groups costs of continuous exposure to injuries. The phylogenetic distance between *E. burchellii* and *M. analis*, which diverged ∼140 million years ago (33), combined with their shared ecological traits, makes this an ideal system for studying the convergent evolution of social wound care and identifying potential new drivers of this behaviour. Together, our empirical and theoretical approaches formalise current arguments for the evolution of social wound care while establishing a baseline for future comparative studies of this behaviour.

## Results

### Costs of raiding pugnacious prey

*E. burchellii* sustains severe injuries during their raids on social insect colonies (**Movie S1**). To quantify the injury rate of raiding parties, we collected over 100 foragers from each of 15 colonies (3995 ants overall). We observed, on average, 11% of injured foragers in a colony (range: 0.6-41%; standard deviation (SD): 11.0, *N*=15 colonies). Injuries were equally likely to occur on the antennae or on each leg pair (**Fig. S1**; Chi^2^ test: χ^2^ =0.12, df=3, p=0.99). The media, the most frequently found body size class during raids (34) (**Fig. S2**), was the most commonly injured ant group among worker classes (**Fig. S1; Table S1;** Dunn test: media-minim p<0.001, media-porter p<0.001, media*-*soldier p<0.001). The other body size classes, typically associated with vertebrate defence [soldiers (31, 35)], transport [porters (34)], or nest tasks [minims (34)], did not show significant differences in their injury rates (Dunn test: soldier*–*minim p=0.79; soldier*-*porter p=0.96; media-porter p=0.97).

### Wound care towards injured ants occurs on-site and inside the nest

To study how injured ants received wound care during the raid, we cut the hind leg of a healthy ant at the femur and released them next to the raiding column. Freshly injured ants remained beside the column and bent their gaster under their body without moving (hereafter referred to as ‘gaster bending’). With the gaster underneath the mesosoma, the ant opened their mandibles and put their antennal scapes together (**Fig. 1a**). Nestmates then walked towards the injured ant and began to allogroom the body. Once an ant found the injury, they licked into the wound with the maxilla and labium for an average of 25% of the time during the first 30 min (**Fig. 1b; Movie S2**). Notably, even though minims make up only 20% of the raid (**Fig. S2**), they conducted 80% of the wound care interactions (62 out of 78 caregivers). In most cases, the injured ant interrupted treatment after five minutes (**Fig. 1c**) and moved a short distance before resuming the gaster bending position for further care. Gaster bending and receiving wound care or allogrooming were strongly associated (**Fig. 1c, Fig. S3**). Healthy individuals also showed gaster bending and allogrooming after being picked up and released (without cutting the leg). While the proportion of individuals bending the gaster and receiving allogrooming did not differ between healthy and injured ants (gaster-bending: Chi^2^ test: χ^2^ =0.03, df=1, p=0.85; allogrooming: Chi^2^ test: χ^2^ =0.14, df=1, p=0.71), the time spent bending the gaster and being allogroomed was significantly higher in the injured (**Fig. 1d, Fig. S4;** bending: mean effects t=4.67, p<0.01; allogrooming: mean effects t=4.33, p<0.01).

**Fig. 1.**
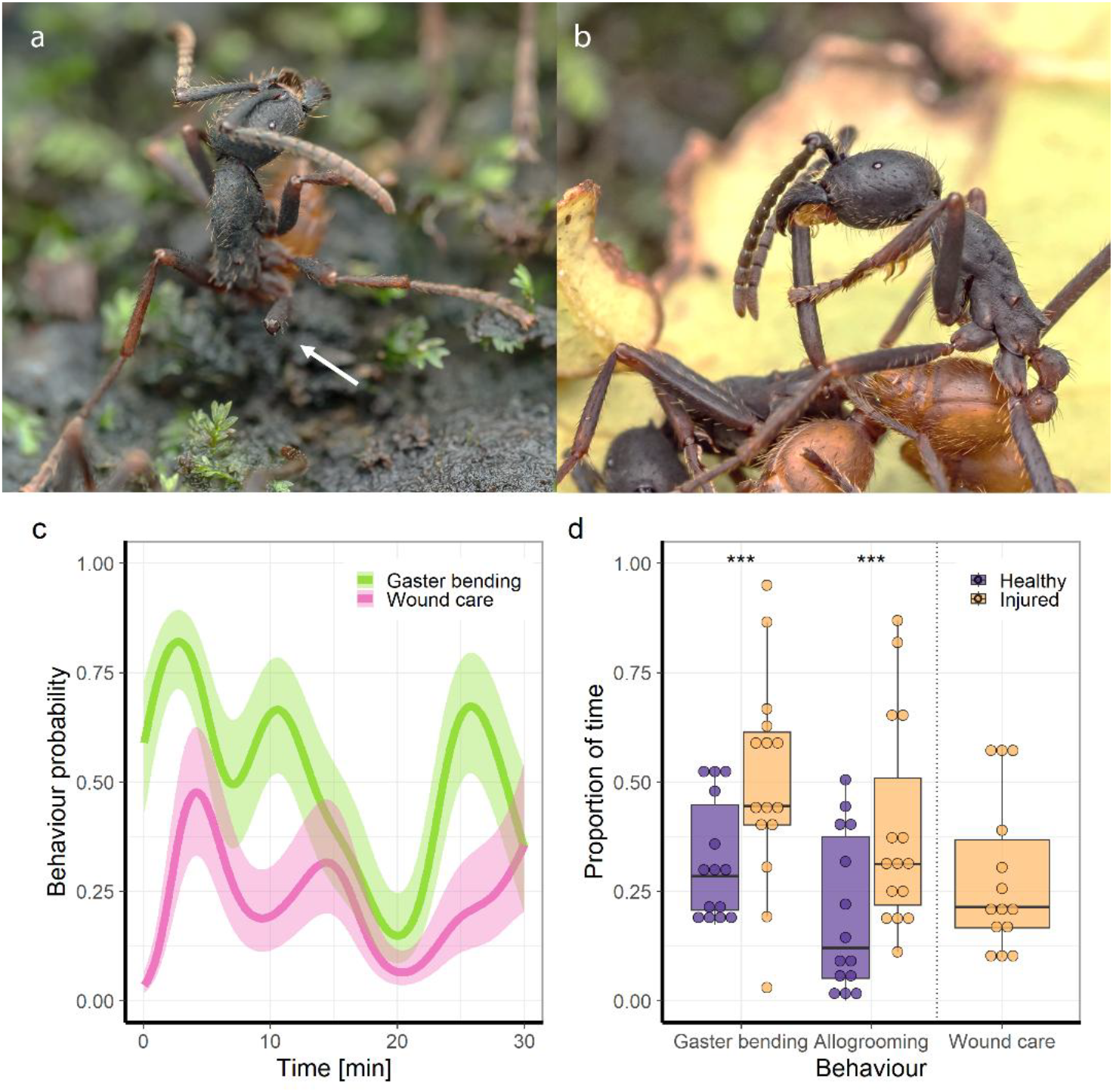
On-site wound care behaviour on injured ants. **(a)** A recently injured ant bends their gaster, spreads their legs, puts their antennal scapes together and opens their mandibles. The white arrow indicates the open wound on the hind right leg. **(b)** Nestmates walk towards the injured ant and groom the body. Once they detect the injury (center), the nestmate licks with the labium and maxilla inside the wound while holding onto the injured limb with the mandibles and legs. Photos in panels a and b were provided by Jeremy Squire. **(c)** Probability of bending the gaster and receiving wound care during the first 30 minutes of getting injured. The presence/absence of the behaviours at intervals of 5 seconds was fitted with a Hierarchical Generalised (Binomial) Additive Model (HGAM). The HGAM model explained 24% of the deviance, with a coefficient of determination *R*^2^=0.29 over 23,650 records within each 5-second interval. The presence of the behaviours, bending the gaster (pink) and wound care (green), was recorded if the behaviour occurred at least once during the interval. The line represents the predicted probability by the model, with the coloured shaded area representing the 95% confidence interval. **(d)** Proportion of time the focal individual spent bending the gaster and opening the mandibles, receiving allogrooming on its body, or on the wound (wound care). 30-minute ethograms compare colour-marked healthy ants (healthy, purple, N=14) to ants with a sterile cut on their hind femur (injured, yellow, N=15). Boxplots depict the first to third quartiles, interquartile range, horizontal lines within boxes are medians, and the whiskers are 1.5 times the interquartile range. ***=p<0.001.

To observe whether wound care behaviour also occurred inside the nest, we created a vertical two-dimensional bivouac (**Fig. S5**). Injured ants whose wounds were exposed to a droplet containing ∼10^5^ bacteria of *Pseudomonas aeruginosa* were released in the nest (infected ants N=65). After release, the infected ants quickly assumed the gaster-bending posture and received allogrooming and wound care inside the bivouac. On four occasions, we observed the minims using their front legs to collect secretions from their metapleural glands and spreading them on the wound with their mouthparts (**Movie S3**).

### Wound care increases the survival of ants facing severe infections

To determine the efficacy of wound care in coping with infections, we compared the survival rates of injured individuals in isolation with or without receiving wound care. Healthy and sterile injured ants had similarly high survival rates (**Fig. 2, Table S2**). While infected injured ants died significantly faster in isolation compared to infected injured ants that were in the nest for the first 12 hours before being placed in isolation (**Fig. 2, Fig. S6, Table S2**). Importantly, infected injured ants that received wound care in the nest showed survival rates similar to those of sterile injured workers (**Fig. 2, Fig. S6, Table S3**). Social wound care effectiveness was also similar among ants treated in their natural bivouac in the field (**Fig. S7**). These results suggest that wound care significantly reduces the mortality from severe infections on the injury. To identify if injured ants participate again in future raids, we quantified injury rates at the start of a raid (before any fighting could occur). Notably, the proportion of injured ants at the start of a raid did not differ from that of an ongoing raid (**Fig. S8**, Kruskal-Wallis; Chi^2^ test: χ^2^ =0.01, df=1, p=0.94), indicating that recovered injured workers participated again in future raids.

**Fig. 2.**
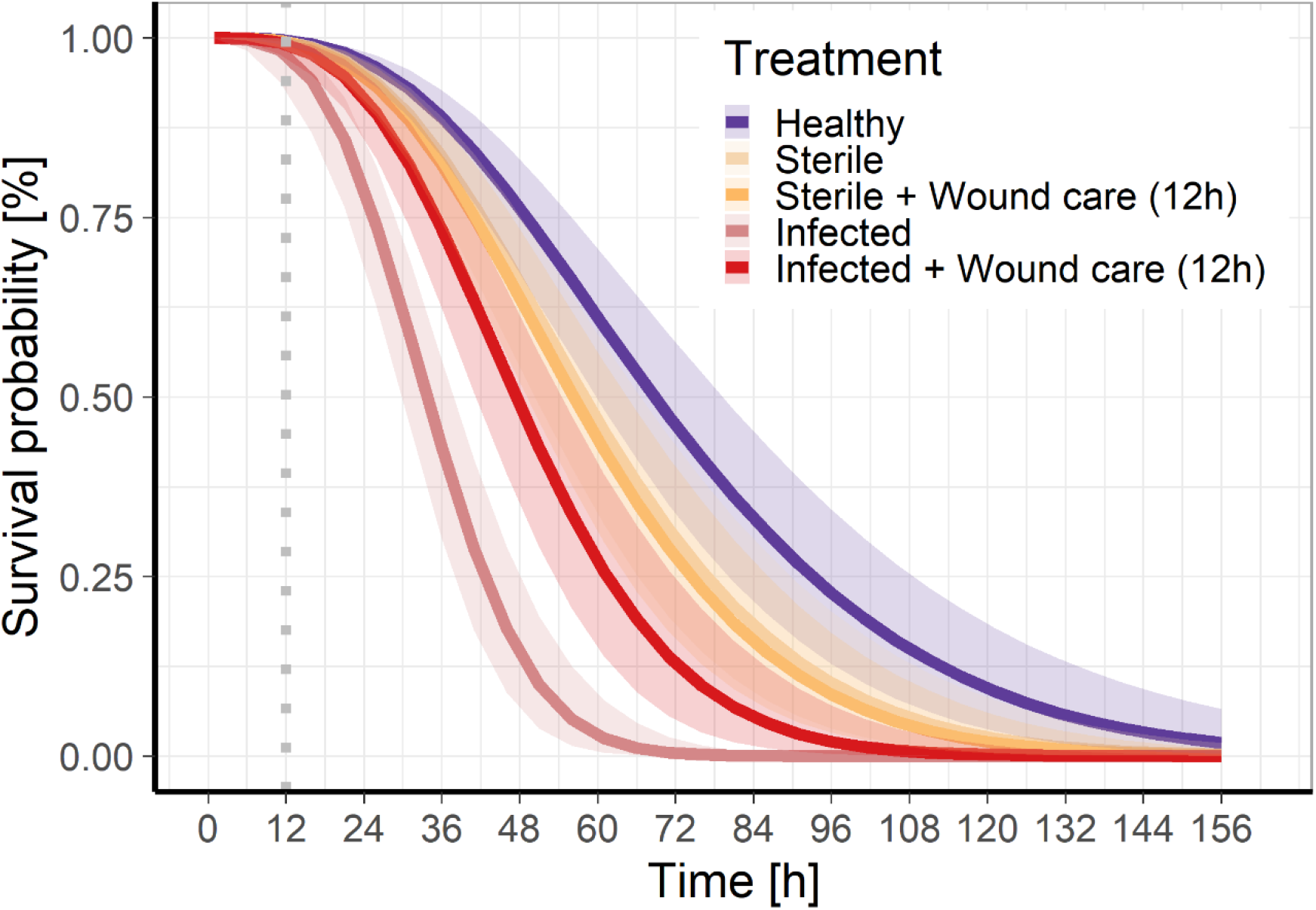
Effect of social wound care on the survival of sterile and infected injured ants. Solid lines show the predicted probabilities, and shaded areas show the 95% confidence intervals from the fitted flexible parametric generalised survival model. Healthy (purple): Non-injured ants; Sterile (yellow): Injured ants with a wound exposed to sterile water; Infected (red): Injured ants with a wound exposed to ∼10^5^ *P. aeruginosa* bacteria diluted in sterile water; Sterile + Wound care (12h) (light yellow) and Injured + Wound care (12h) (light red) denote injured ants that were colour marked, and allowed to receive care in the nest for the first 12 hours. The dotted grey line indicates the moment the sterile and infected ants were recaptured and placed in isolation from the laboratory nest (**Fig. S5**). All workers were kept in sterile isolation chambers on sterile soil. Data combined from two colonies: Healthy (N=41), Sterile (N=40), Sterile + Wound care (N=41), Infected (N=41), Infected + Wound care (N=40). Details of statistical analysis are provided in **Fig. S6** and **Table S2**. Social wound care effectiveness was also similar among ants that received treatment in their natural bivouac (**Fig. S7**).

### CHC cues for injury condition

To test whether ants use cuticular hydrocarbons (CHCs) as cues to identify injury condition, we performed sterile and infected injuries on ants in isolation and quantified the changes in CHCs directly after injury (control), five hours, and sixteen hours later (**Fig. 3**). Overall, we found 41 CHC compounds on the ants (**Table S4**), with variations mainly driven by intraspecific colony differences (**Fig. S9, Table S5;** Colony effect PERMANOVA: F=86.22, df=2, R^2^=0.38, p<0.001). While injury condition and time after injury also showed significant differences in the CHC composition, the variance they explained was low (**Fig. S10, Table S5;** Group effect PERMANOVA: F=5.45, df=3, R^2^=0.04, p<0.001; time: F=7.42, df=1, R^2^=0.02, p<0.001). Pairwise comparisons showed no clear discrimination among healthy, sterile, and infected injured ants over time (**Table S6**). To exclude the possibility of a social effect on the communication pathway, we also kept ants in a sub-colony, where we observed similar patterns to those of ants kept in isolation (**Fig. S11, S12, Table S7, Table S8**).

**Fig. 3.**
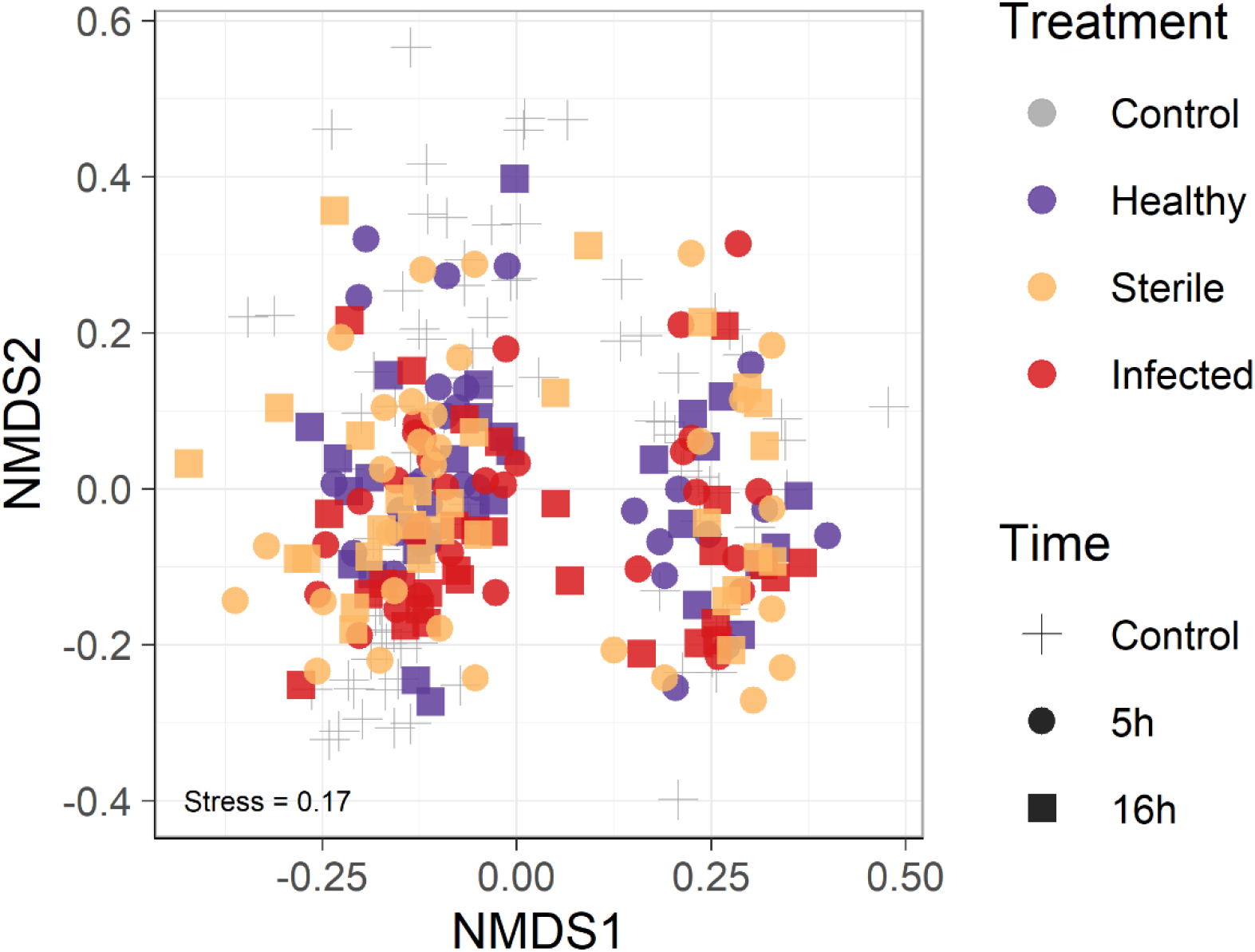
CHC profiles for ants that received infected or sterile wounds over time. CHC profile similarities are displayed in bi-dimensional Non-metric Multidimensional Scaling (NMDS) based on a dissimilarity matrix from Bray-Curtis distances. Colours represent ants kept in isolation that received infected or sterile injuries whose CHCs were extracted at different times. Control (grey) corresponds to ants that were either healthy or received infected or sterile injuries, and their CHC extraction occurred immediately after manipulation. Shapes represent the time of CHC extraction (hours) after injury. Control (N=87), Healthy 5h (N=30), Sterile 5h (N=30), Infected 5h (N=30), Healthy 16h (N=29), Sterile 16h (N=30), Infected 16h (N=30). Treatments did not differ in their CHC compositions but showed significant differentiation by colony. Inside each colony, the profiles show a stronger differentiation over time, rather than due to injury or infection (**Fig. S10, Table S6**). CHC’s also remained similar across sterile and infected groups kept in isolation or inside artificial sub-colonies (**Fig. S11, Fig. S12, Table S7, Table S8**).

### Injury rate, not group size, determines the benefits of social wound care

To study how injury rate and colony size influence the evolution of social wound care, we developed a theoretical model in which we consider wound care as a helping behaviour that reduces the added mortality from injuries **(Fig. 4a and b)**. Our model confirms that the maximum colony growth rate (the eco-evolutionarily relevant parameter for reproduction, reflecting the fastest colony fission with the largest number of ants) decreases as colonies are under higher injury rate regimes **(Fig. 4c)**. But, if injured ants can recover with the help of nestmates, colony growth rate increases as the percentage of helped ants increases, up to levels comparable to those in colonies without injuries **(Fig. 4c)**. Importantly, colony size does not affect the benefits of wound care on colony growth rate, because both the injured and helped fraction scale directly with colony size **(Fig. S13)**.

**Fig. 4.**
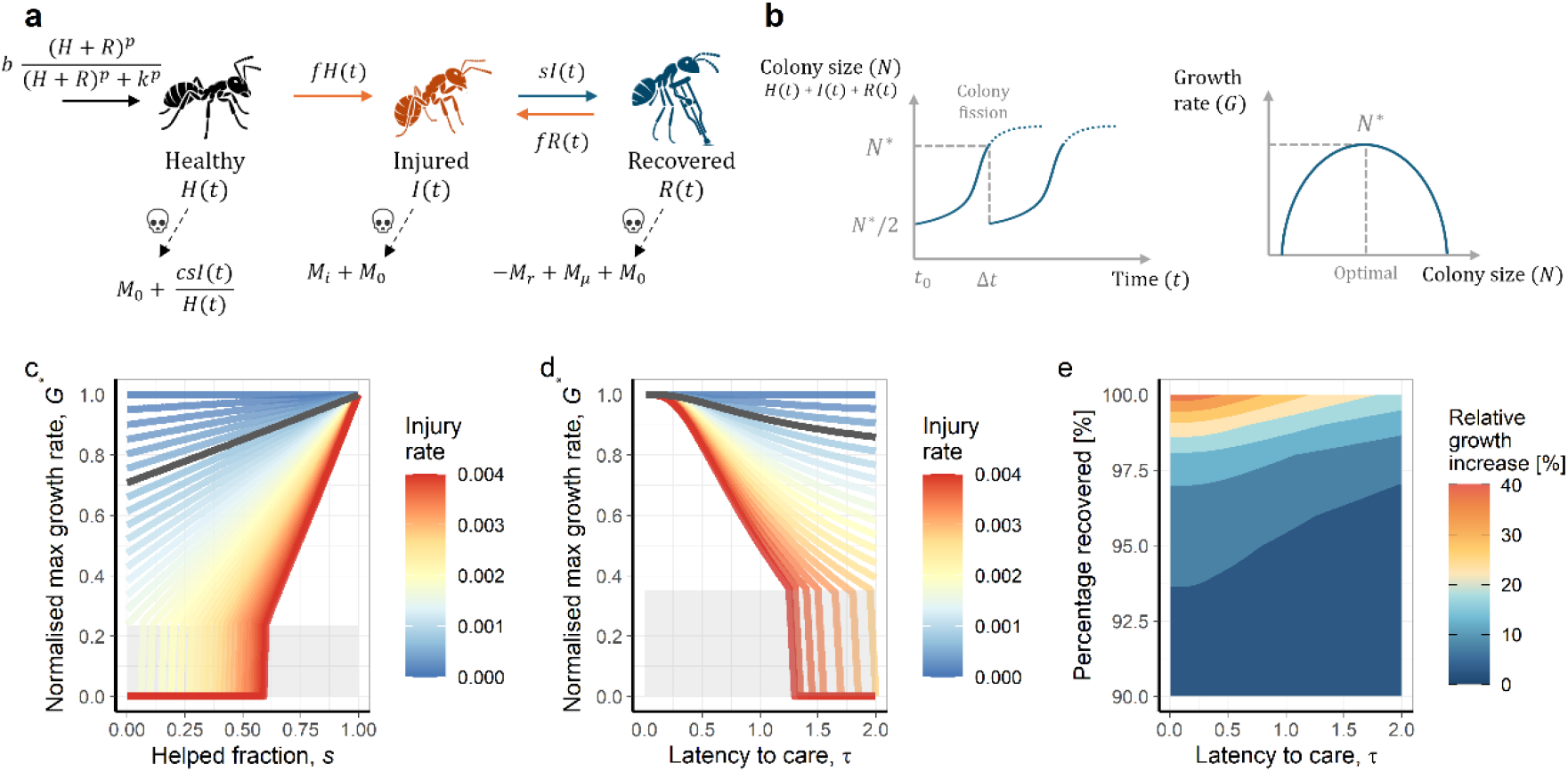
Framework for the role of injury rate, care timing and recovery in shaping colony growth. **(a)** Colony-growth dynamics. The colony produces a maximum *b* of new ants per time point, which is density-dependent on the current number of healthy (*H*), and rescued (*R*) ants. The density-dependence strength of *b* is shaped by the scale parameter *k* and shape parameter *p*. All ants have a per capita mortality *m*_0_. In addition, healthy ants suffer a per capita injury rate *f*, increasing the base mortality by *m*_*i*_ . A fraction of ants is helped *s* every time step, reducing the increased mortality of the injury by *m*_*r*_. Recovered ants can experience new injuries at the same rate *f*. If costs *c*, are assumed, they are incurred by healthy ants that provide wound care on injured ants. **(b)** Colony-reproduction dynamics. The number of ants in a colony increases until the colony reaches an optimal colony size *N*^*∗*^ in which the colony reaches the highest growth rate to split at a size *N*^*∗*^/2. Note that the shape of the function determining the optimal colony size changes with the parameter combination. **(c)** Effect of injury rates and the helped fraction on colony-level dynamics represented by the difference model. The maximum colony growth rate *G*^*∗*^ represents the ratio between the optimal colony size for splitting *N*^*\**^ and the time when colony growth is the highest. The helped fraction *s* represents the fraction of injured ants that received wound care. The black line illustrates the growth rate dynamics for *E. burchellii* at the observed mean injury ratio (11%), with the colour gradient representing the empirically observed range of injury rates across colonies. The shaded grey area denotes the approximate minimum helped fraction required to increase colony growth rate. **(d)** Interaction effect of injury rates and latency to care on colony growth in the ODE model. Quick care provides accelerating returns on colony growth, especially at high injury regimes. The latency to help τ represents the time an injured ant waits until receiving help with each τ unit representing one day. The black line illustrates the growth rate dynamics for *E. burchellii* at the observed mean injury ratio (11%), with the colour gradient representing the empirically observed range of injury rates across colonies. The shaded grey area denotes the approximate minimum required latency to care to increase colony growth rate. The effects of wound care costs are shown in **Fig. S14a. (e)** The joint effect of injury recovery and latency to care on colony growth in the ODE model. High injury recovery provides stronger benefits for colony growth than shorter care latencies. Colours represent changes in the relative growth rate in bins of five units. The effects of wound care costs are shown in **Fig. S14b**. Parameter details are in **Table 1**.

To understand the importance of the latency and efficacy of social wound care, we modified the model such that τ represents the average number of time steps an injured ant must wait before receiving care (hereafter, latency to care). Our model emphasises that quicker first response times accelerate colony growth, suggesting strong drivers to reduce waiting time **(Fig. 4d)**. In our case, the longer it takes for an injured ant to receive care, the greater the chance that pathogens infect the wound or that an infection progresses. In addition to the importance of receiving quick wound care, a high injury recovery is mandatory for substantial increases in colony growth rate **(Fig. 4e)**. Notably, the benefits of wound care only manifest above certain thresholds **(Fig. 4c, d, and e)**. When the helped fraction or injury recovery rate fall below their respective minima, or when latency to care exceeds a critical maximum, wound care becomes insufficient to compensate for colony losses at high injury rates, thus preventing net colony growth. These results underscore the importance of receiving early and efficient wound care for the evolution and retention of social wound care behaviours.

**Table 1.** Parameter values used for finding numerical solutions.

| parameter | value | meaning | source |
| --- | --- | --- | --- |
| $b$ | 1700 | maximum number of ants produced per day | (46) |
| $k$ | 200,000 | density dependence scale of birth rate | fit with data from (46) |
| $p$ | 4 | density dependence shape of birth rate | fit with data from (46) |
| $\theta$ | 100 | colony size scaling factor to interpolate between *E. burchellii* and *M. analis* | |
| $m$ | 600,000 | colony size at the moment of reproduction | |
| $f$ | 0.0006 | per capita injury rate | tuned to fit a splitting time of 1 year and injured percentage of ~11% |
| $c$ | 0-0.5 | per capita added mortality on healthy ants providing care | |
| $m_0$ | 0.0025-0.003 | base per capita ant mortality | tuned to fit a splitting time of 1 year and injured percentage of ~11% |
| $m_i$ | 0.7 | per capita added injury mortality | fit using data in Fig. 2 |
| $m_r$ | 0-$m_i$ | per capita reduced mortality by receiving help | |
| $n_1$ | 5 | shape factor of base survival curve (continuous model only) | fit using data in Fig. 2 |
| $n_2$ | 7 | shape factor of survival curve from injury (continuous model only) | fit using data in Fig 2. |
| $s$ | 0-1 | fraction of helped ants (discrete model) | |
| $\tau$ | 0-5 | average time an ant waits until receiving help (continuous model) | |

## Discussion

In this study, we connect social wound care in an army ant with its proximate and ultimate benefits in coping with risky predator-prey interactions. Army ants suffer from open wounds during their raids, but effectively handle lethal wound infections by performing social wound care. This care is partitioned into two phases: (1) an on-site wound care at the raiding site after receiving a fresh injury, and (2) a follow-up wound care treatment which can involve the use of the metapleural gland secretions in the bivouac. CHCs, a well-known source of communication in social insects, do not change after injury and likely do not play a central role in signalling injury or infection in *E. burchellii*. Contrary to the current hypothesis that social wound care behaviours have greater fitness benefits in small societies (26, 36), our theoretical model shows that the benefits of care are independent of colony size.

Army ants constantly battle with other ants to acquire prey (35). Fights in *E. burchellii* occur with several arboreal *Camponotus* species (37, 38), at the cost of suffering open wounds. Open wounds that are not properly attended to can develop into infections, threatening the individual’s survival (10, 17). We detected that, on average, 11% of foragers, ∼100,000 ants in a colony, carry at least one injury, injuries that, when infected, the innate immune system alone is insufficient to resolve. Social wound care thus effectively reduced mortality caused by infected wounds, as also observed in other ant species (10, 26). This suggests that without wound care, colony size would be dramatically reduced, with our models estimating that the colony growth rate could be 32% lower without it.

Treatment timing on open wounds is a key axis for individual survival (15, 39, 40). We show that injured army ants not only receive treatment at the nest, but, contrary to previous studies in ants, social wound care is also conducted immediately after injury in the field. Injured ants at the raiding ground display a stereotypical sequence of behaviours, from bending their gaster to opening their mandibles, until attracting their nestmates. This on-site care could be a response to the long raid times in army ants, often lasting over 12 hours (35). By the time the injured ants would return to the bivouac, an infection could have already festered in the wound (10, 15). Thus, by receiving on-site care, ants likely remove contaminants from the wound and reduce pathogen load (41–43), thereby extending the time window for further successful treatments in the nest. This on-site care may therefore be a strategy unique to species with prolonged foraging times. Other collective hunters, such as *M. analis*, conduct short raids on termite foraging sites that rarely exceed 30 minutes before returning to their nests (44). The time between injury and treatment is thus seldom longer than 15 minutes (45), and likely does not necessitate on-site wound care (15).

The moment of injury relative to the raiding phase could also shape caregiver availability and the latency of their care, thereby influencing treatment effectiveness. Worker availability could be particularly important since on-site wound care was primarily performed by minims, which are far less common in raids than the frequently injured medias (46). Despite their small size imposing greater energy costs for long-distance travel, minim participation in raids is thought to directly benefit colony fitness by plugging potholes and building living bridges, thereby increasing traffic flow and food retrieval (47, 48). Here we identify an additional benefit of minim inclusion in raids: the provision of on-site care for injured nestmates. These cost-benefit dynamics suggest that minims may represent an emerging caste-level specialisation in injury care.

The high genetic relatedness and density in the nest of social insects highlight the importance of rapid detection and targeted treatment of infected nestmates (49–52). Cuticular hydrocarbons (CHC) are a reliable source of communication (53), and non-surprisingly, their change follows immune challenges across several taxa (54). Our analyses of CHC profiles show that CHC changes occurred only over time, and not as a response to wounding or infection. These patterns were consistent regardless of whether nestmates were present, suggesting that social isolation does not mask the communication of injury or infection via CHC changes. At the same time, an adaptive signalling pathway should not only increase the chances of the diseased ant’s survival (55, 56), which is achieved when the ant is treated in the nest, but also reduce the odds of diseases spreading to kin (50, 57, 58). If the wound care treatment elicited during the gaster-bending posture at the raiding site and inside the bivouac is sufficient to cope with wound infections, selection for a new signalling pathway could be redundant. Therefore, specific signalling of injury condition through CHCs may not be necessary.

### Evolution of social wound care

Social wound care is part of the collective disease defences that social groups display to cope with pathogen-induced mortality and disease outbreaks (10, 15, 17, 51, 59). However, unlike other social immunity related behaviours, which primarily prevent or reduce disease risks at the level of the colony (59, 60), social wound care distinguishes itself as a directed response towards injured individuals. Injury creates a temporary window during which opportunistic pathogens, otherwise harmless to healthy individuals (10, 41, 61), can exploit the compromised integument and establish lethal systemic infections. Social wound care behaviours, such as amputations (15, 16) and antimicrobial care (10) act within this window to reduce pathogen load and promote wound healing; preventing infections from becoming systemic (15) or providing therapeutic care for already established infections (10). These behaviours extend the injured individual’s lifespan, thereby maintaining workforce size and thus benefiting colony fitness.

Evolutionary explanations for social wound care have highlighted group size as a potential driver (26, 62). Frank and Linsenmair (2017) propose that the benefits of helping an injured individual increase as group size decreases, since the relative value of each individual is greater in smaller groups. In small groups, the loss of a single individual can more directly compromise collective success as argued for early human groups (63) or cooperative breeders such as birds (64). Further comparative studies could shed light on the causal relationship between group size and other social traits in the evolution of social wound care. However, our model demonstrates that the benefits from caring for injured ants are independent of group size. The fraction of cared individuals scales with colony size – the more ants are in a colony, the more can suffer injuries, meaning the benefits to reproductive rate depend on injury and care rates rather than absolute colony size. Our model further highlights an additional fitness consequence of care: each recovered worker accelerates colony growth, thereby advancing the timing of colony reproduction. This effect is particularly pronounced in fission-reproducing species, such as army ants like *E. burchellii*, but it extends to any species in which colony growth influences reproductive output.

Other factors, such as injury frequency and lethality, have also been suggested to influence the evolution of these behaviours (26). Our model formalises and disentangles their interaction with other factors, such as the degree, latency, and efficiency of caring. As expected, high injury rates and lethality translate into slower colony growth, which can be buffered by high degrees of efficient care. Therefore, selection could favour the evolution of different wound care behaviours that synergistically cope with the costs associated with high frequency and lethality of injuries. Lastly, the elapsed time until receiving care was fundamental in increasing colony growth rate. This could explain why joint action of social wound care in two phases, on-site and inside the nest, may provide accelerating returns to the colony’s reproductive rate in *E. burchellii*. In army ants, in which obtaining food is costly, social wound care can buffer these costs and create an eco-evolutionary feedback loop of increasingly risky fights with greater food gains. The high benefits of social wound care on colony growth, without apparent cost, suggest that these behaviours may be much more widespread across taxa facing risky interactions.

## Conclusion

Our study considers social wound care as a crucial helping behaviour that enhances individual survival and fitness in a social group. Social wound care takes two phases, an on-site care at the raiding site which may act as a prophylactic strategy, and wound care inside the nest with potential use of metapleural gland secretions to combat infections. These behaviours thereby reduce the costs associated with risky predator-prey interactions during extensive raids. We further provide a theoretical framework that formalises the current hypothesis on the evolution of social wound care. We show that group size is not a fundamental axis in shaping the evolution or retention of this behaviour. Instead, the interaction between injury rate and lethality shapes the costs of risky interactions that are buffered by social wound care. The strong accelerating returns in the colony’s reproductive rate from caring for the injured suggest that these behaviours may be much more widespread than previously thought.

## Materials and Methods

The on-site wound care behaviours were recorded by annotating 30 min ethograms of experimentally injured ants at the raiding site. The efficacy of social wound care was assessed by measuring the survival of isolated infected ants with the opportunistic pathogen *P. aeruginosa* after receiving wound care in artificial nests. We investigated changes in CHCs over time by extracting sterile and infected injured ants in hexane, followed by gas chromatography/mass spectrometry analyses. To understand how helping behaviours, such as social wound care, translate into fitness, we formalised a discrete-time model accounting for within- and colony-level dynamics. We further transformed the model into a continuous model to understand the relevance of latency to care, injury rate and recovery on fitness. Detailed methods and protocols for the injury rate survey, ethograms, pathogen culture, wound care efficiency, infection signalling, and theoretical models are provided in ***SI Appendix***.

## Data, Materials, and Software Availability

All study data are included in the article and/or ***SI Appendix***. Simulation code and data analysis scripts are available under https://doi.org/10.5061/dryad.jdfn2z3qf (65).

## Acknowledgments

We thank the Organization for Tropical Studies (OTS), and the Staff at La Selva Research Station, especially Orlando Vargas, Enrique Castro, Danilo Brenes and Genesis Brenes, for their valuable support during the fieldwork. We thank Ana Abarca, Pia Wapler, and Jakob Jansen for their field assistance. Daniel Kronauer, Christoph von Beeren and Jonathan Saragosti for their advice on the laboratory nest set-up. The workshop at the Biocentre of the University of Würzburg for the nest construction. Special thanks to Daniel Rodríguez-León for his meaningful discussions and advice on the chemical analyses. Doris Waffler for her technical assistance and Zsolt Kárpáti for his help on the CHC samples derivatisation. Gaurav Athreya for his discussions on the theoretical model. Special thanks to Hanna Haring and Jeremy Squire for their documentation of army ant behaviour through scientific illustrations (Hanna) and photography (Jeremy). Juan J. Lagos-Oviedo and Erik T. Frank are supported by the DFG Emmy Noether Program #511474012 and the Hector Fellow Academy. This study was performed under the permission of the Comisión Nacional para la Gestión de la Biodiversidad (CONAGEBIO) R-015-2023-OT-CONAGEBIO and R-027-2025-OT-CONAGEBIO.

