## Supplementary Material for "Social wound care dynamics in an army ant society"

### Supporting Information Methods

#### Study site and organism

This study was conducted in a lowland tropical forest located in La Selva Biological Station (10°25′19″N - 84°00′54″W) in Costa Rica. Observations and experiments were conducted from March to April, October to December 2023, and February to March 2026. We studied the army ant *Eciton burchellii foreli* Mayr, 1886, a top tropical predator of arboreal ants, social wasps and other arthropods (1–3). Colonies range from 500.000 to 2 million individuals and reproduce by fission (4, 5). Army ants are a classic example of mass-raiding behaviour, with hundreds of thousands of workers typically raiding from dawn until sunset (6, 7). Additionally, colonies alternate between a migratory and a stationary phase. During the migratory phase, the colony raids to feed their larvae during the day and migrates every evening for approximately 25 days. In the stationary phase, the colony stays in the same location, raiding at lower intensity for approximately 20 days (5, 6, 8). We located colonies by following their foraging trails back to the bivouac.

Workers of *E. burchellii* exhibit a continuous body size variation within the most abundant worker class, in addition to two bigger, distinct classes: porter and soldier, sometimes referred to as submajors and majors (9). The porter class has remarkably bigger whitish heads with elongated legs (10), whereas the soldier class also has distinctly hook-shaped elongated mandibles (9, 10). We subdivided the most common worker class into two groups based on their maximum head width (9); minim (0.96-1.49 mm) and media (1.50-1.80 mm). Individuals were visually classified into these body size classes for behavioural observations and experiments.

#### Survey of injury rate

To identify the proportion of injured ants per body size class in a colony, we sampled at least a hundred ants from all group sizes in a raiding column in which ants were already carrying back prey. Fifteen raiding events corresponding to different colonies were collected for a total of 3995 sampled individuals. An injury was considered the partial or complete loss of a limb or antennae in an ant. We recorded the frequency of these injuries across body size groups. Because different numbers of ants were sampled in each raid, affecting the estimations of injured ants, we confirmed a non-association between the number of collected ants and the proportion of injured ants (Pearson correlation test: $S=734$, $r=-0.31$, $p=0.26$). To determine the proportion of injured ants carrying injuries from a previous raid, we collected and recorded the injury ratio from colonies when leaving the nest, before a fight could occur. Ants were collected during the first hour of the raid for each colony, when all raiding ants were leaving the nest. Data was collected for six colonies for a total of 1557 observed ants.

**Wound care behaviour description**

To study the behavioural repertoire of injured ants compared to healthy individuals, we carried out on-site ethograms. Foraging ants were taken from a raiding column at least two hours after the raid started. Each ant was painted on their mesosoma with a red Acrylic Edding® marker and then left to dry in a box for 10 minutes. Afterwards, the focal individual was placed four cm from the raiding column and at least five metres away from the bivouac. For the artificially injured ants, we cut the right hind leg with sterilised scissors at the centre of the femur, whereas healthy ants did not receive any injury. During the ethograms three behaviours were quantified. (1) Gaster bending: the focal individual was bending their gaster and opening their mandibles. (2) Allogrooming: the focal individual was touched by nestmates with their mouth parts. (3) Wound care: nestmates licking directly into the injury. Ethograms were conducted for 30 minutes directly after placing the focal ant back into the raiding column. Observations in which the focal individual was lost within the first five minutes were discarded. A total of 15 ants from four colonies were observed for both healthy and injured groups (N=15).

**Pathogen culture and infection**

To perform controlled wound infections, we injured and infected the open wound on the hind right leg with *Pseudomonas aeruginosa*, a generalist pathogen commonly found in soil. The isolated strain previously obtained by Frank et al. 2023 (11) was cultured in Tryptic Soy Agar medium at 26°C. Open wounds were exposed to 20 µL drops of distilled water during 2 seconds, containing approximately $10^5$ bacteria, corresponding to a concentration of 0.05 optical density (OD), as done in previous studies (11, 12). OD was measured with an Ultrospec 10 Cell Density Meter. To ensure consistent pathogen activity, we used bacteria in the growth phase, between 12 and 24 hours after replating.

**Wound care inside a bivouac-like environment**

To observe wound care behaviour in a bivouac-like environment, we created an artificial vertical nest of acrylic plates and cork (**Fig. S5**). Three thousand ants carrying some larvae were collected from a migration column and placed inside the nest overnight. Ants created a bivouac-like structure by creating agglomerations with ants and larvae. To determine whether ants provide wound care to infected ants in the artificial setup, as observed at the raiding site, we infected a total of 65 ants using the method as in the “Pathogen culture and infection” section. Injured ants were introduced to the artificial nest one at a time. Despite the artificial nest's advantages in reproducing bivouac-like behaviours, quantifying ethograms over prolonged periods of time was impossible. Thus, we followed the ants whenever they were visible and qualitatively recorded the wound care behaviours. Two types of behaviours were observed. (1) Wound care: the injured ant bends their gaster, while receiving allogrooming and wound care by nestmates. (2) Antimicrobial wound care: nestmates collected their metapleural gland secretions with their front legs and applied them to the wound. Observations were done for 17 hours in three sub-colonies (N=15; N=20; N=30 released injured workers per colony). Metapleural gland treatment was recorded with a Nikon L310 camera.

**Efficacy of wound care behaviour on ant survival**

To determine the effectiveness of social wound care in the nest on the recovery of ants with infected injuries, we compared the survival of ants treated in the laboratory nest with that of ants without care. We created two sub-colonies by separately collecting ~3,000 workers with larvae from two migratory colonies (**Fig. S5**). To allow the ants to habituate to the new environment, the sub-colonies remained undisturbed overnight before the experiments. We compared the survival of five groups: (1) healthy (N=41): ants without any visual injury, (2) sterile (N=40): ants that were injured with a sterile pair of scissors whose wounds were exposed to a 20 µL sterile distilled water solution, (3): infected (N=41): ants that received the same sterile treatment but whose wounds were instead exposed to a pathogen solution (see “Pathogen culture and infection” for further details), (4) sterile + treated 12 hours (N=41), and (5) infected + treated 12 hours (N=40) who received care in the artificial nest for 12 hours before being retrieved and put in isolation. All ants were previously painted with Acrylic Edding® marker for identification.

All ants were kept in individual isolation chambers, consisting of cylindrical glass tubes with a 3 cm diameter and 5 cm height. These containers were filled with 2 cm of surface soil, covered with aluminium foil and autoclaved at 130°C for 40 minutes. Humidity was maintained by moistening the soil with 1 mL of sterilised water at the beginning of the trial. No food was provided during the assay. The isolation chambers were kept at a mean environmental temperature of 28°C during the experiment. Ant survival was quantified for the first 160 hours.

We further confirm the effectiveness of wound care inside the nest under natural conditions on the recovery of infected ants by comparing the survival of infected ants that received care in the bivouac with those kept in isolation. In this case, we compared four groups: (1) Healthy, (2) sterile, (3) infected, and (4) infected + treated, which were retrieved from their colony 36 h after release. To recapture infected individuals released in the colony, we collected and painted the mesosoma with a red Acrylic Edding® marker on 100 ants from a foraging column. We infected and released the marked ants at the end of the day (19:00h) at a 1 m distance from the bivouac (see above for infection procedure). Ants stayed for 36 hours in the bivouac, during which the colony did not forage because of strong rainfalls. Ants were collected at the beginning of the following foraging day until the first 20 marked ants were obtained. A total of 20 ants per group were placed in isolation chambers (N=20). Ant survival was quantified for the first 160 hours.

**Use of CHCs for injury and infection detection**

To investigate whether the CHC profile of ants facing an infection changed over time, we collected 150 ants from the return column and treated them as described in the section above on ant survival. (1) Healthy: uninjured ants, (2) sterile: ants that received a sterile injury, or (3) infected: ants whose wound was infected with *P. aeruginosa* at 0.05 OD. We extracted the CHC profiles for all ant groups at 0, 5, and 16 hours after treatment (**Table S4**). Ants treated for five and sixteen hours were kept in sterile isolation chambers, as described above. Ten replicates were performed for each time and group, for a total of 90 individuals per colony (N=10 per group). We repeated the experiment for three different colonies to account for intracolonial variation in the initial relative abundances of CHCs. To confirm whether ant mortality was due to infection, we ran the experiment in parallel to additional survival experiments (as described above) and compared the survival curves between the groups. Infections significantly increased mortality compared with healthy or sterile-injured ants (**Fig. S15, Table S10**). We used between 11 and 15 ants per group across 3 colonies for the parallel survival assays (**Fig. S15**). To study the effect of the environment on CHC changes across treatments, we compared the CHC profiles of ants that remained in the artificial bivouac (social environment) with those in isolation (isolation environment) within a single colony at zero (control), five, and 16 hours after treatment with another parallel survival experiment confirming the efficacy of infections (**Fig. S16**). For the survival experiment, we used healthy (N=13), sterile injured (N=13), and infected ants (N=13).

**Chemical extractions and analyses of CHC**

To study how CHCs changed over the course of the infection, we performed extractions by submerging ants in 1 mL of *n*-hexane for 10 minutes and then evaporating the solution to approximately 100 µL. We used 1 µL extracts for gas chromatography/mass spectrometry (GC-MS) analyses, performed on a gas chromatograph (6890) coupled to a mass-selective detector (5975) from Agilent Technologies. The GC was equipped with a DB-5 capillary column (0.25 mm inside diameter × 30 m; film thickness, 0.25 µm; J&W Scientific). Helium was used as the carrier gas, with a constant flow rate of 1 mL/min. A temperature program from 60°C to 300°C with

5°C/min and finally 10 min at 300°C was used, with data collection starting 4 minutes after injection to exclude the solvent peak. The mass spectra were recorded in the electron ionisation mode, with an ionisation voltage of 70eV and a source temperature of 230°C. To calculate the relative abundance of CHC compounds, we calculated the area under each compound peak using the integration function of MassHunter (Agilent Technologies) and divided each peak by the total area under all CHC peaks. We aligned CHC peaks across samples by using *GCalignR* v.1.0.5 (13).

We analysed the fragmentation pattern acquired from the MS to identify CHCs. Double bond positions of unsaturated hydrocarbons were identified by obtaining dimethyl disulfide derivatives as done in (14). We removed all non-hydrocarbon compounds from the analysis (polar compounds and contaminants). Similarly, we removed compounds that were present in less than half the samples in all groups or in only trace amounts (<0.1%) across all samples. Due to failed extractions or high concentrations of contamination, we excluded four samples from the experiment in isolation and six from the one including the social environment. We compared the relative abundance of each compound and compound class (Alkanes, Methyl-branched alkanes, Alkenes, and Alkadienes) across the different treatments.

**Theoretical model for the evolution of wound care**

To understand the interaction dynamics and the importance of injury rate and colony size on the evolution of social wound care (15, 16), we formalised a theoretical model that accounts for the individual benefits and colony reproductive consequences of this behaviour. We construct a model that considers the within-colony dynamics of birth, death, injury, and help, and the colony-level dynamics of growth and fission. Parameter values, their explanation and sources are provided in **Table 1**.

**Individual-level / within-colony dynamics**

To understand how these helping behaviours contribute to colony reproduction, we developed a discrete model in which each time step represents one day. From one day to the next, new ants are produced, a fraction of the ants experience injuries, and a fraction of them are helped. We assume that colonies are composed of clonal ants whose growth and dynamics are described by a system of difference equations based on a compartment model (**Fig. 4a**). The discrete within-colony dynamics from one day to the next are described by the following difference equations:

$$\Delta H = b\frac{(H+R)^p}{(H+R)^p + k^p} - fH - m_0 H$$

$$I = f(R+H)$$

$$\Delta R = sI - fR - (m_0 + m_i - m_r)R$$

Where ants can be either healthy (*H*), injured (*I*), or recovered from the injury (*R*). New healthy ants are produced according to a density-dependent birth rate, with a maximum birth rate $b$, based on the current number of healthy and recovered ants (*H*+*R*). The density-dependence is controlled by parameters $k$ and $p$, which determine the scale and shape of the relationship respectively. At low colony sizes, the birth rate shows accelerating effects, and diminishing returns at large colony sizes. Thus, these parameters create a natural soft threshold, reflecting that a minimum colony size is required for colony maintenance.

The healthy ants have a baseline mortality $m_0$. Healthy and recovered ants can get injured during foraging at a constant daily injury rate $f$. The injured ants are helped depending on the colony-level phenotype of helping $s$, which represents the fraction of injured individuals that are helped within a day. Following helping, a recovered ant has, in general, a higher mortality compared to healthy ants, represented by $m_0 + m_i - m_r$. An injured ant must be helped by the end of the day, otherwise it succumbs (*e.g.* infections). Therefore, at the beginning of the next day, the number of healthy and recovered ants changes depending on the dynamics of the previous day, but the number of injured ants is reset to 0.

To understand the importance of these helping behaviours across taxa that differ in colony size, we included the parameter $\theta$, which scales colony size by scaling birth rate $b$, and the strength of the density-dependent growth $k$ **(Fig. S13).** Thus, the scaling factor $\theta$=1 represents species with small colony size (~4000 ants) and has *b*=17*, k*=2000, while $\theta$=100 represents an army ant colony (~400,000 ants), with *b*=1700*, k*=200,000.

The above model sheds light on the importance of the fraction of injured ants helped for a myriad of injury rates and colony sizes. However, the timing at which an ant receives help may be relevant in determining individual survival and colony growth. We studied this by converting our discrete-time model into a continuous-time system of differential equations. In this model, the number of healthy, injured and recovered ants changes continuously over time. The rates of birth, injury, mortality, and helping are now interpreted as instantaneous rates rather than fractions per day. Now, injured ants have an elevated mortality rate given by the base mortality $m_0$ and that imposed by the injury $m_i$. In addition to the continuous nature, we also capture more precisely the survival curves of **Fig. 2**. That is, the inverse S-shaped curve of gradual decrease, followed by a rapid decrease and then a gradual decrease in survival probability. This requires introducing multiple stages for each mortality process (base ($n_1$ stages) and injured-induced ($n_2$ stages)). Injured ants are helped depending on the colony-level phenotype of helping rate *s,* representing the instantaneous fraction of injured ants that are helped in a unit of time. Thus, $\tau = 1/s$ is the average time that an injured ant must wait before it is helped and is the quantity we use for analysis. Whether or not an ant is helped depends on the interplay between the injury-induced mortality and the latency to care. If the added mortality is very high, the ant may die before receiving help. Once helped, injured ants recover, and their mortality is reduced by an amount $m_r$. A full recovery means that the mortality of the recovered individuals returns to the base mortality. Healthy ants that help injured individuals incur a cost in the form of a mortality probability due to the act of caring. The continuous-time within-colony dynamics are described by the following differential equations:

$$g(x) = \frac{x^p}{x^p + k^p};\ \lambda_0 = n_1 m_0; \lambda_c = n_1 cs \frac{\sum_{j,l} I_{j,l}}{\sum_j H_j}; \lambda_i = n_2 m_i; \lambda_r = n_2(m_i - m_r)$$

$$\frac{dH_1}{dt} = bg(H+R) - (\lambda_0 + \lambda_c + f)H_1$$

$$\frac{dH_j}{dt} = (\lambda_0 + \lambda_c)H_{j-1} - (\lambda_0 + \lambda_c + f)H_j \quad j = 2, \dots, n_1$$

$$F_j = f \cdot \left( H_j + \sum_{l=1}^{n_2} R_{j,l} \right)$$

$$\frac{dI_{j,l}}{dt} = F_j \mathbb{1}_{[l=1]} + \lambda_0\, I_{j-1,l} \mathbb{1}_{[j>1]} + \lambda_i\, I_{j,l-1}\, \mathbb{1}_{[l>1]} - (\lambda_0 + \lambda_i + s)I_{j,l}$$

$$G_j = s \sum_{l=1}^{n_2} I_{j,l}$$

$$\frac{dR_{j,l}}{dt} = G_j \mathbb{1}_{[l=1]} + \lambda_0 R_{j-1,l} \mathbb{1}_{[j>1]} + \lambda_r R_{j,l-1} \mathbb{1}_{[l>1]} - (\lambda_0 + \lambda_r + f)R_{j,l}$$

Here, the healthy compartment of individuals $H$, has $n_1$ stages for the base mortality process. The index $j$ represents the stages of this base mortality process. The base mortality and injury-induced mortality are two independent mortality processes and thus are tracked independently. That is, when a healthy ant is injured and subsequently recovered, its stage in the base morality process continues and is not reset. Therefore, for each stage in the base mortality process, we need $n_2$ stages in the injury-induced mortality process, resulting in $n_1 \times n_2$ stages for the injured $I$ and recovered $R$ compartments. The index $l$ represents the stage in the injury-induced mortality process. The $\lambda$s in the equations are shorthand for the rates of leaving from the different stages.

$F_j$ represents the total rate of ants in the $j$th stage of the base mortality process that are getting injured. $G_j$ represents the total rate of injured ants in the $j$th stage of the base mortality process who are helped. The indicator function $\mathbb{1}_{[...]}$ is 1 if the condition within the square brackets is true and 0 otherwise. Ants of all healthy stages can help injured ants, and thus, the cost of caring for injured individuals is proportionally shared across all stages of the healthy compartment.

**Colony-level dynamics**

The previously described within-colony dynamics determine how quickly the colony can expand in size and the maximum size it can achieve. While the colony-level dynamics describe the reproduction of whole colonies themselves, through growth and fission, and the maximisation of the overall growth rate over evolutionary time. Consider that a colony splits when it reaches a size *N*, into two colonies of equal size $N/2$. This colony size at splitting is something that the colony can *choose* in addition to the other parameters. For a given colony size, we assume that the colony composition reaches equilibrium. That is, the composition of the colony, the fraction of the different compartments and stages, remains the same at each fission event. At this equilibrium, the time a colony takes to grow from $N/2$ to *N* is termed as the splitting time $\Delta T$. The eco-evolutionarily important quantity is the overall colony growth rate $G = N/\Delta T$. The splitting time and the overall colony growth rate depend non-linearly on the colony size. On the one hand, a large colony size results in a longer splitting time, leading to a lower overall growth rate. On the other hand, a small colony size translates into fast splitting time, but also results in low overall growth rate, due to the small size. Therefore, there is an intermediate colony size $N^*$ which maximises the overall growth rate. For this optimal colony size $N^*$, the splitting time is $\Delta T^*$, and this produces the maximum overall growth rate $G^* = N^*/\Delta T^*$. Over evolutionary time, the population will settle into this optimal colony size for a given parameter combination. We then compare the optimal overall colony growth rate across parameters.

**Model parameterisation and simulations**

Colony size in *E. burchellii* ranges from 300,000 up to 2 million ants (4, 6, 17). The birth rate or rate of production of ants varies across colony sizes. We use data from (18) to fit a Hill function

$$g(x) = \frac{x^p}{x^p + k^p}$$

using least squares and obtain the density dependence scale $k$=194,919 $\pm$ 5210 and density dependence shape $p = 4.28 \pm 0.48$ parameters. For computational ease and convenience, we use $k$=200,000 and $p$=4.

Injury-induced mortality may vary depending on the type of prey attacked or pathogens infecting the wound (19–22). For parameterising the mortality processes, we used data from **Fig. 2** and fitted it with the Erlang survival function (23)

$$S(t) = e^{-nmt} \sum_{j=0}^{n-1} \frac{(nmt)^j}{j!}$$

to obtain the mortality $m_i = 0.688 \pm 0.02$ and the number of stages for the healthy $n_1 = 5$ and injured $n_2 = 7$ compartments. Since the survival measurements for these fits were done in an experiment in isolation, not reflecting the lifespan of a healthy ant, we do not use it to parametrise the base mortality $m_0$.

The parameters of base mortality $m_0$ and injury rate $f$ were tuned to obtain a colony at optimal equilibrium, which displays ~11% of injured and recovered ants in the colony (**Fig. S1**) and a colony splitting time of ~1 year, as suggested by natural history observations for an army ant colony (6, 17, 18). This gives $m_0 = 0.003$ and $f$=0.0006. To capture the observed variation in the injured fraction of foragers between 0 and 47%, we vary the injury rate from 0 to 0.004. The other parameters are varied over a wide range to examine the system's properties.

**Statistical analyses**

We used a significance level of 0.05 as reference for statistically significant p-values. All analyses were performed using the R language v.4.5.2 (R Core Team 2023) with R Studio v.2023.06.0. environment (RStudio Team 2020).

**Wound care behaviour**

To test whether a specific group of ants exhibited a different proportion of injuries, we performed a Kruskal-Wallis test with a posterior Dunn *post hoc* test. To compare the absolute difference between gaster bending and allogrooming between healthy and injured individuals, we calculated the proportion of time each behaviour occurred for each individual. We fitted an ordinary regression model for the additive effects of behaviour and group. We posteriorly predicted the mean pairwise differences in behaviours between healthy and injured individuals using the package *emmeans* v.2.0.1 (24).

To analyse the temporal patterns of wound care, gaster bending and allogrooming behaviours in healthy and injured ants, we modelled the presence or absence of these behaviours in five-second intervals through Hierarchical Generalized Additive Models (HGAMs). In all HGAMs we included the interaction of behaviour with time as a fixed effect while accounting for the colony as a group structure. We estimated independent smooths at each level of the behaviours by including the individual and their colony as nested random intercepts. Analyses were performed using the package *mgcv* v.1.9.3 (25). Model check was performed by using *DHARMa* v.0.4.7 (26).

To quantify the efficacy of wound care behaviour on the survival of infected individuals in the laboratory nest, we conducted Generalised Parametric Survival Models using the package *rstpm2* v.1.7.1 (27, 28). We modelled the survival probability using a stratified model with smooths for the baseline Hazard for the group treatment and time in the nest. Colony ID was included as strata. Three internal knots were used for the splines after comparing models varying the number of knots and time varying effects. Likelihood-ratio tests were used to select the best model. We further compared the survival of infected ants in the natural bivouac by conducting a Cox proportional Hazard regression model using the package *survminer* v.0.5.2 (29) We modelled worker survival probability according to group treatments. The survival probabilities were illustrated using Kaplan-Meier cumulative survival curves.

**Chemical analyses of infections**

To assess how CHC profiles change when ants sustain an injury or an infection, we calculated the Bray-Curtis dissimilarity indices between samples of healthy, sterile injured, and infected injured ants across time zero, five, and 16 hours after injury. We performed a multivariate analysis of variance (PERMANOVA) with 9999 permutations to assess differences between treatments over time with the package *vegan* v.2.7-2 (30). Since CHC profiles across treatments were similar at time zero (PERMANOVA: n=86, F= 0.98, $r^2$=0.02, p=0.11), we merged these groups into one, hereafter called control. To investigate the influence of the environment (either social or in isolation) on the CHC profile change upon injury and infection, we compared the Bray-Curtis dissimilarity indices across time zero (control), five and 16 hours after injury in ants kept inside the laboratory nest, or in isolation. Comparisons across groups were performed using PERMANOVA, including the interaction between environment, treatment, and time.

CHC profiles across treatments and time points were visualised using a non-metric multidimensional scaling (NMDS) ordination plot. To determine specific group differences in CHC composition, we performed pairwise comparisons across groups using Adonis tests. We controlled for multiple-comparison false discovery rates by applying the Benjamini-Hochberg correction to p-values.

To study how the relative abundances of compound groups in the CHC profiles changed for ants facing an infection, we fitted Generalised Linear Mixed Models (GLM) on the relative percentages in a CHC profile for each compound group depending on treatment and time. Colony ID was added as a random factor, and the model was fitted using *lme4* v1.1.38 (31), before calculating the variance table for an ANOVA type II. We predicted the mean pairwise differences in relative percentages using the *emmeans* package v.2.0.1 (24).

**Model simulations**

We ran simulations in Julia v1.10 with the *DifferentialEquations.jl* v7.15 package, using a custom script, which is provided as part of the data and code (32).

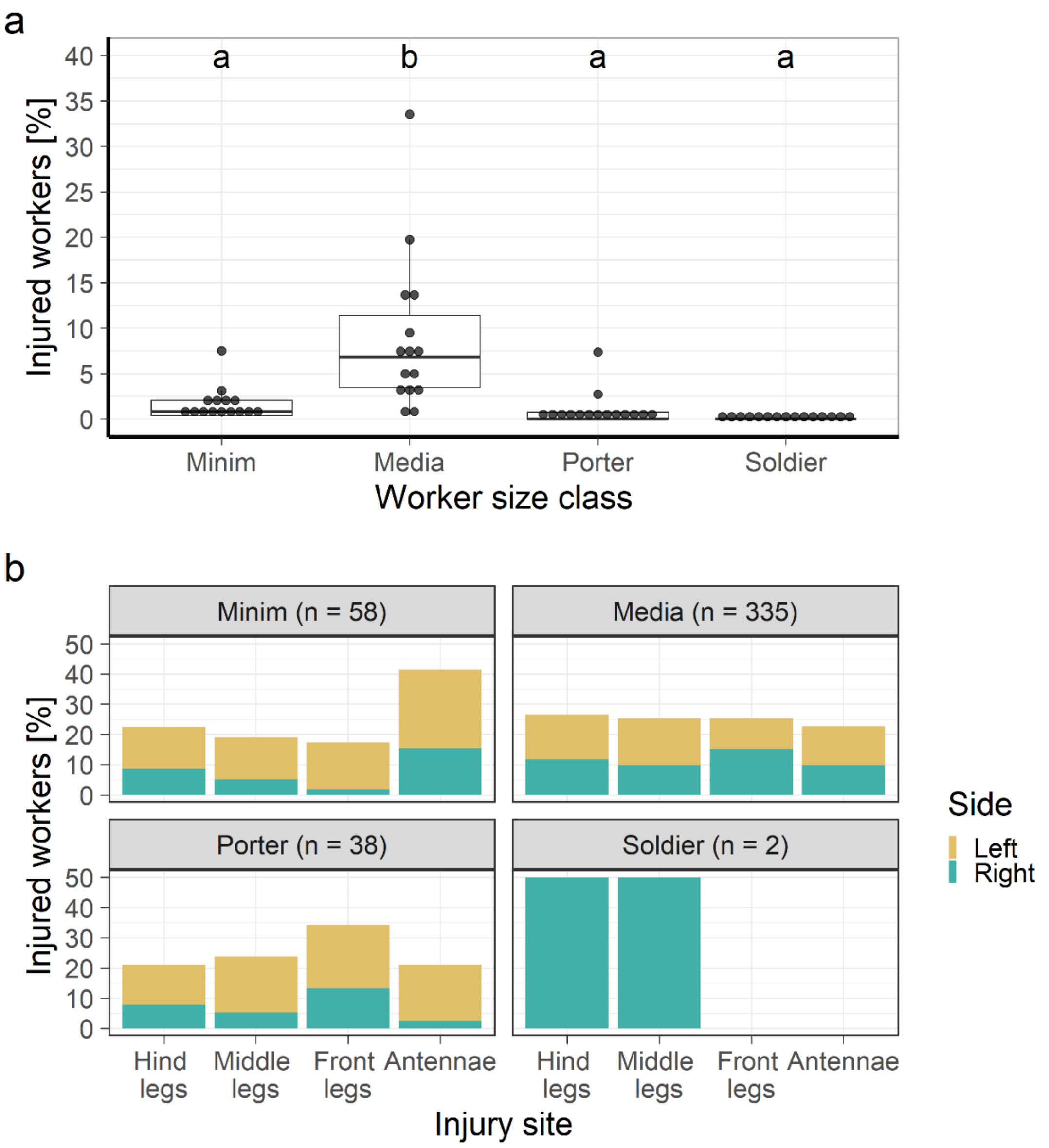


**Fig. S1. Distribution of injuries among body size classes. (a)** Percentage of injured ants per body size class. Size classes are arranged in ascending order by their body size. Body size classes showed significant differences in their injury rates (Kruskal-Wallis: $\chi^2$=39.55, df=3, p<0.001. Each point corresponds to the mean injury ratio calculated for each raid. The mean injury ratio for *E. burchellii* (11%) was obtained by averaging the mean injury ratio values across 15 raids. Boxplots depict the first to third interquartile range, horizontal lines within boxes are medians, and the whiskers are 1.5 times the interquartile range. Different letters denote statistically significant differences in the Dunn *post hoc* test **(Table S1)**. **(b)** Detailed distribution of the injuries per body size class across all raids. Pooled data on injuries revealed that all injuries are evenly distributed across the four body regions (Chi² test: $\chi^2$=0.12, df=3, p=0.99), and that there is no side bias for the media class (Chi² test: $\chi^2$=1.58, df=1, p=0.21). Due to the low sample size, the other groups were not statistically considered. Injury distributions were obtained from observations in 15 raids (for a total of 433 injured workers out of 3995).

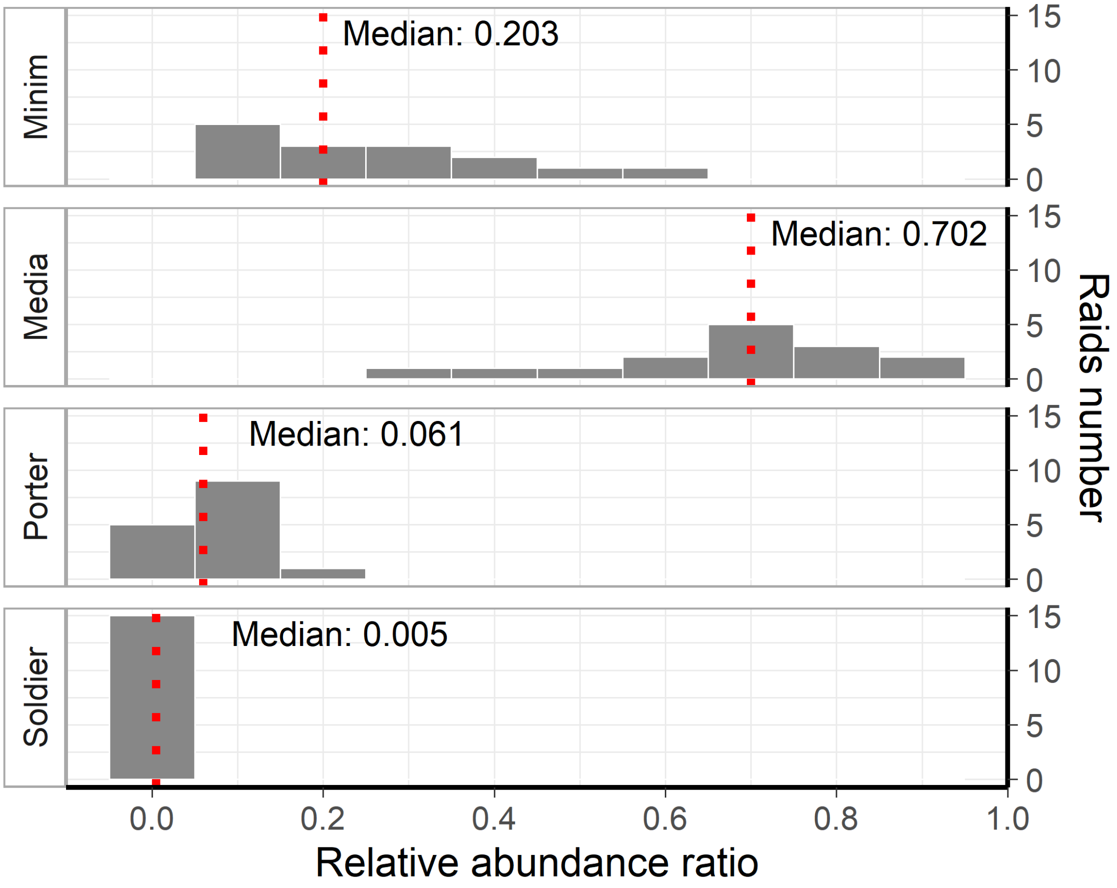


**Fig. S2. Relative abundances for each body size class in the raids of *E. burchellii*.** Histograms represent the relative abundance of body size classes estimated for each studied raid (N=15). Each bin has a width of 0.1 in the relative abundance ratio. The dotted red line represents the median value for each class.

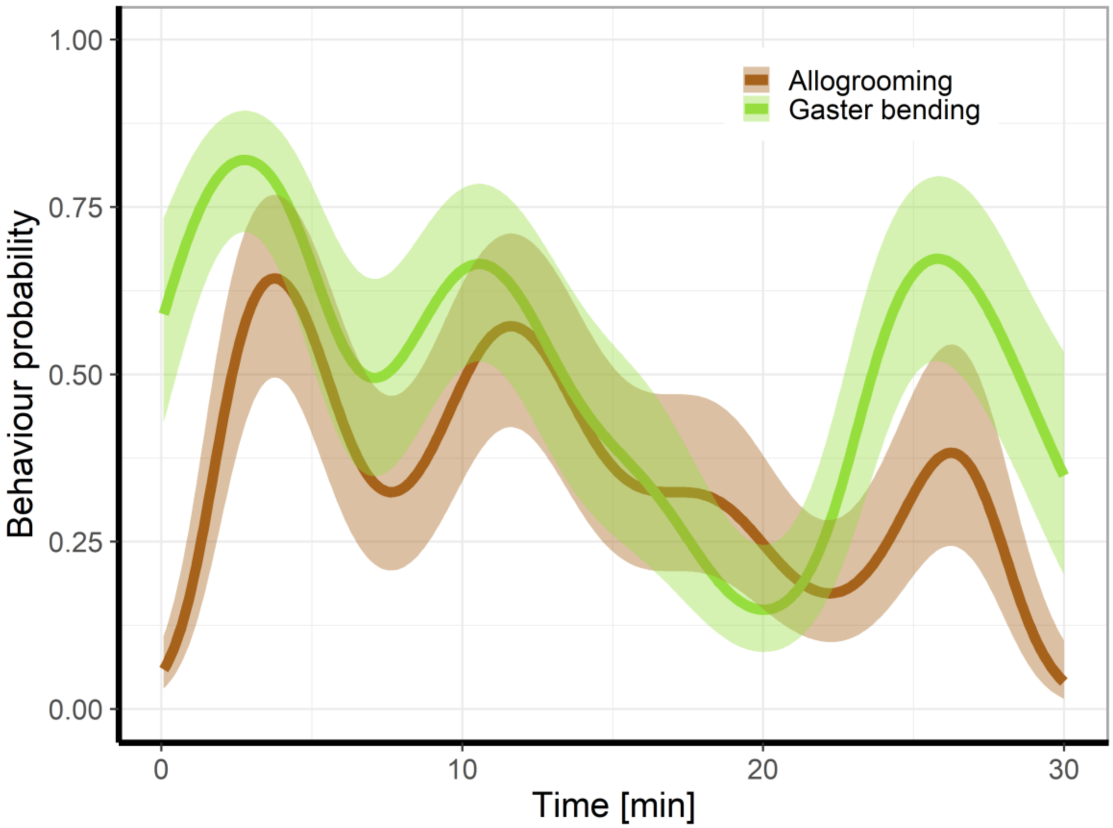


**Fig. S3. Conditional probabilities for the association between allogrooming and gaster-bending in injured workers over 30 minutes.** The presence/absence of the behaviours in intervals of 5 seconds was fitted with a Hierarchical Generalised (Binomial) Additive Model (HGAM). The presence of the behaviour was recorded if the behaviours (gaster-bending, green) and (allogrooming, brown) occurred at least once during the interval. The line represents the predicted probability by the HGAM, with the coloured shaded area representing the 95% confidence interval. The individual response within a colony contributed to explaining the variance ($p<0.001$, e.d.f.=14.40, $\chi^2=3783.50$). The interaction between time and treatment (smooth term) was significant for allogrooming ($p<0.001$, e.d.f.=8.85, $\chi^2=550.40$) and gaster-beding behaviours ($p<0.001$, e.d.f.=8.81, $\chi^2=891.80$). HGAM explained 24% of deviance, with a coefficient of determination $R^2=0.29$ over 23,650 observations. N=14 ants for three colonies.

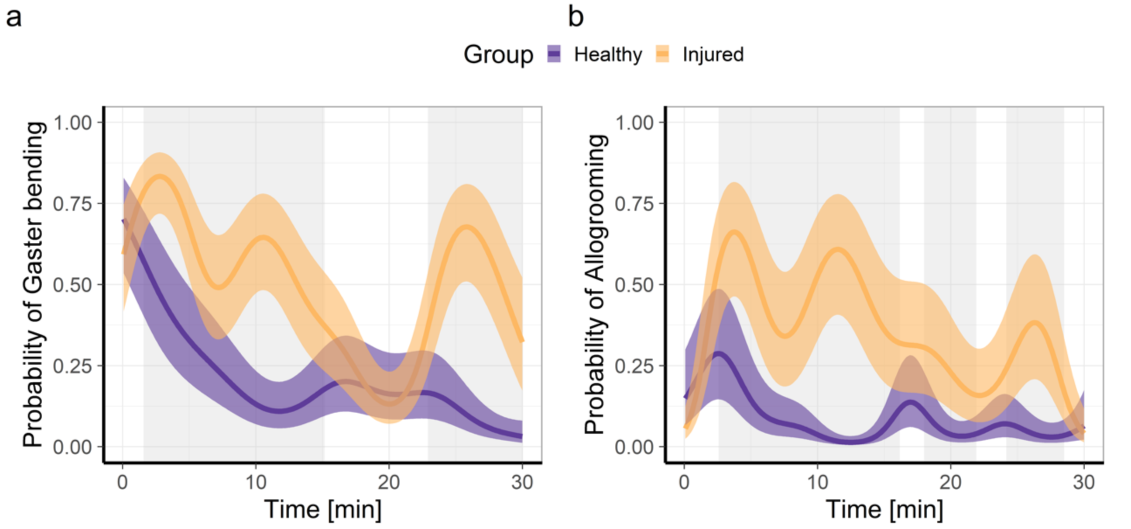


**Fig. S4. Conditional probabilities for the proportion of time spent bending the gaster and receiving allogrooming for healthy and injured individuals during raids.** The presence/absence of the behaviours was quantified in intervals of 5 seconds and fitted with a HGAM. The line represents the predicted probability by the HGAM, with the coloured shaded area representing the 95% confidence interval. Grey shaded bars indicated periods in which the probability of the behaviour between healthy (purple) and injured (yellow) ants was significantly different ($p<0.05$). Each group contains information of 14 individuals from three colonies. **(a)** Time in the gaster bending position over 30 minutes for healthy and injured ants. Injured ants were 32% more likely to receive allogrooming than healthy ants (z-value=2.90, $p<0.01$). The individual response inside a colony contributed to explaining the variance ($p<0.001$, e.d.f.=26.70, $\chi^2$=1950.50). The interaction between time and ant condition (smooth term) was significant for healthy ($p<0.001$, e.d.f.=7.37, $\chi^2$=495.30) and injured conditions ($p<0.001$, e.d.f.=8.81, $\chi^2$=755.70). HGAM explained 24% of deviance, with a coefficient of determination $R^2$=0.30 over 13,057 observations. **(b)** Time receiving allogrooming over 30 minutes for healthy and injured workers. Injured workers were 31% more likely to receive allogrooming than healthy workers (z-value=3.27, $p < 0.01$). The individual response within a colony contributed to explaining the variance ($p < 0.001$, e.d.f.=26.48, $\chi^2$=2354.40). The interaction between time and ant condition (smooth term) was significant for healthy ($p<0.001$, e.d.f.=8.98, $\chi^2$=419.30) and injured conditions ($p < 0.001$, e.d.f.=8.86, $\chi^2$=560.80). HGAM explained 27% of the deviance, with a coefficient of determination $R^2$=0.32 over 13,057 observations.

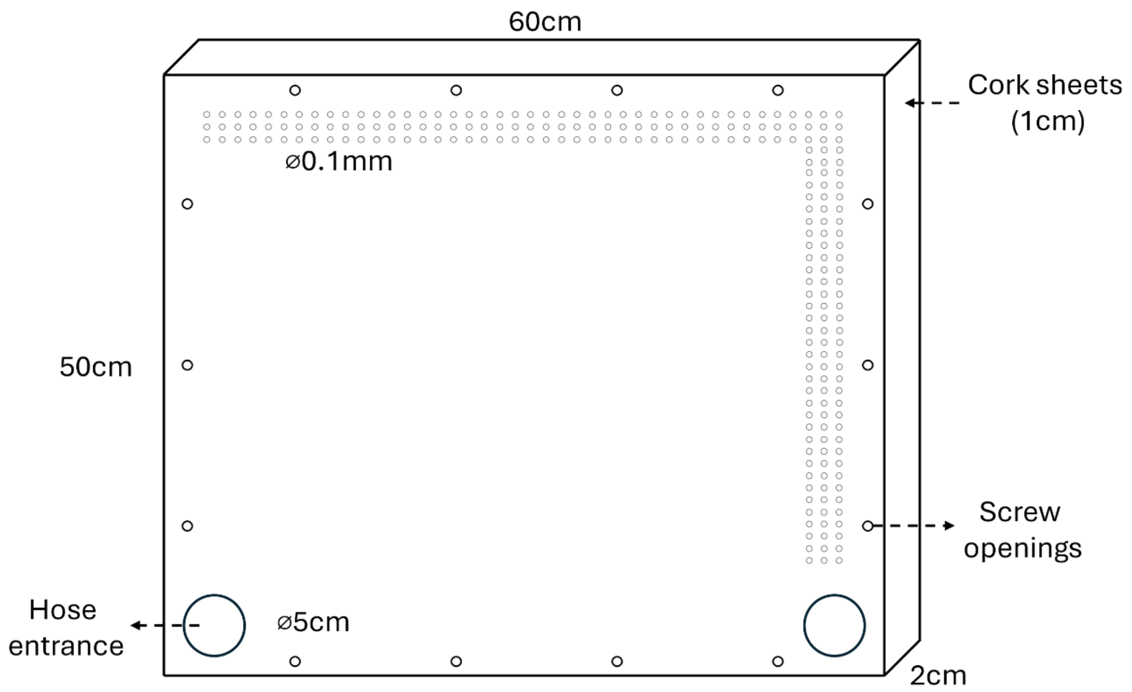


**Fig. S5. Artificial nest setup.** The artificial nest was constructed with transparent acrylic plates and cork sheets, connected by plastic screws. Small holes along the edges of the acrylic plate permitted airflow through the cork material. Two hose entrances are located opposite each other on the lower edge of one plate. One 30-cm hose was connected to a box with soil collected near the bivouac, to which we added grasshoppers. For the observation of wound care under bivouac-like conditions, we collected approximately 2000 workers carrying larvae from a migration column. After a few hours, workers formed chambers where groups of larvae were kept.

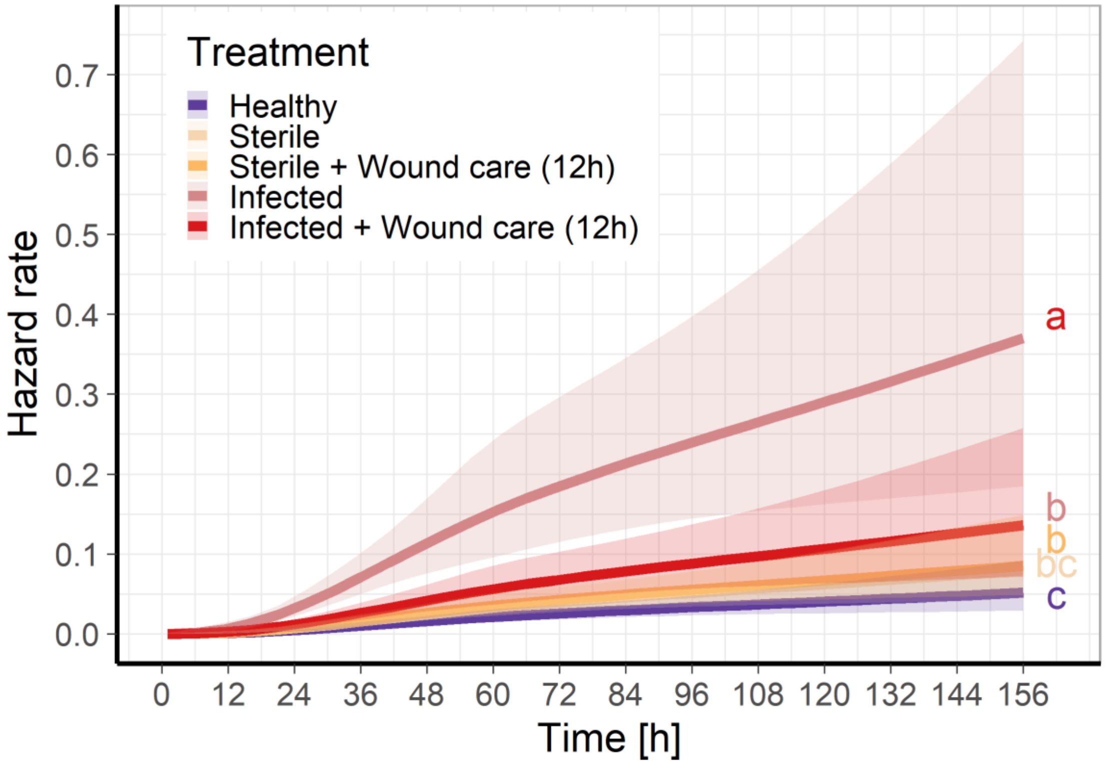


**Fig. S6. Instantaneous hazard rates for the survival of sterile or infected injured ants with or without receiving wound care**. Healthy (purple): Non-injured ants; Sterile (yellow): Injured ants with a wound exposed to sterile water; Infected (red): Injured ants with a wound exposed to ~$10^5$ *P. aeruginosa* bacteria diluted in sterile water; Sterile + Wound care (12h) (light yellow) and Infected + Wound care (12h) (light red) denote sterile and infected injured ants that were colour marked, and allowed to receive care in the nest for the first 12 hours before being placed in isolation. Different letters depict statistically significant differences in the estimated marginal means for the linear predictors from the flexible parametric generalised survival model (**Table S2**).

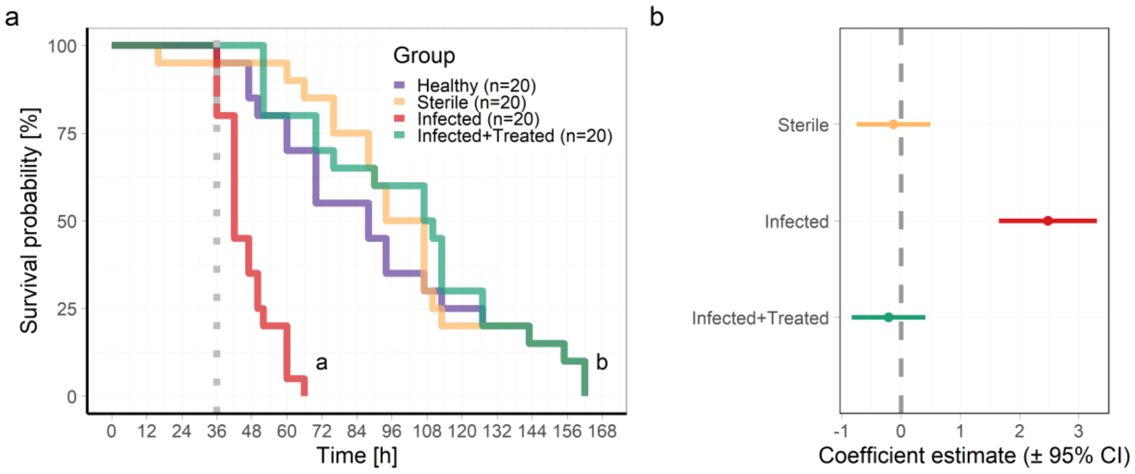


**Fig. S7. Effect of wound care in the natural bivouac on the survival of injured ants**. **(a)** Kaplan-Meier cumulative survival rates show the survival probability of ants in different conditions. Healthy (purple): Non-injured ants; Sterile (yellow): Injured ants with a wound exposed to sterile water; Infected (red): Injured ants with a wound exposed to ~$10^5$ *P. aeruginosa* bacteria diluted in sterile water; Infected+Treated (green): Infected ants that were colour-marked and allowed to receive wound care in the field for the first 36 hours. The dotted grey line shows the moment the infected ants were recaptured and placed in isolation when leaving the nest. No ant left the bivouac for the first 36 hours due to heavy rains. Workers were kept in sterile isolation chambers on sterile soil. Different letters depict statistically significant differences in the Cox proportional hazards regression model (**Table S3**). **(b)** The forest plot shows the estimated hazard ratios for the survival of infected and treated ants. Ants without any injury (healthy ants) were used as a reference group. Coefficient estimates of the Hazard ratios are on a logarithmic scale. Coefficients exponentially transformed are shown in **Table S3.**

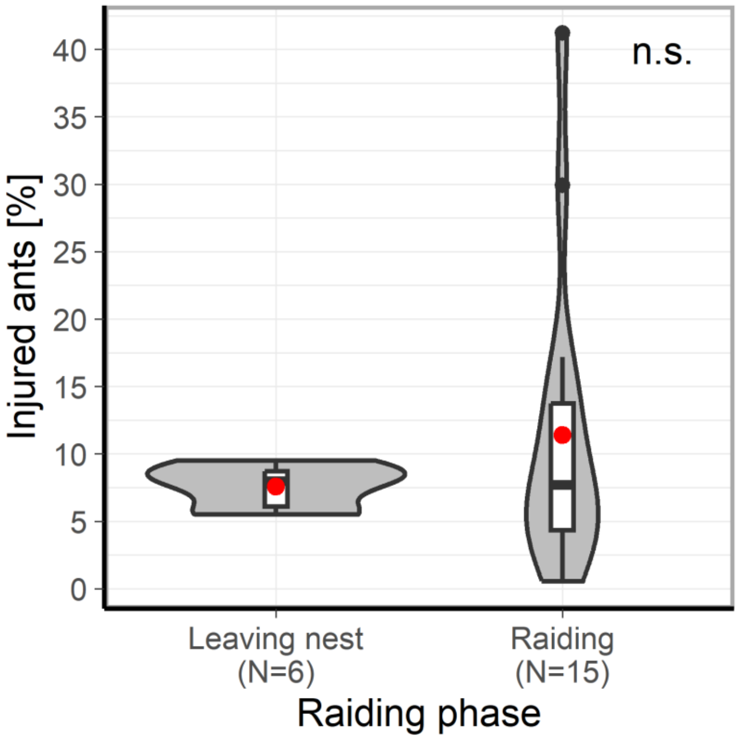


**Fig. S8. Relative abundance of injured ants depending on the raiding phase.** The proportion of injured ants was similar regardless of whether ants were leaving for the raid or during raiding (Kruskal-Wallis; Chi$^2$ test: $\chi^2$ =0.01, df=1, p=0.94). Red dot represents the mean injury ratio for each group (leaving ants: 7.6%, raiding ants: 11.4%). Boxplots depict the first to third quartile of the interquartile range, horizontal lines within boxes are medians, and the whiskers are 1.5 interquartile range. Kernel densities are shown as shaded areas in violin plots. Each data point represents the proportion of injured ants per raid.

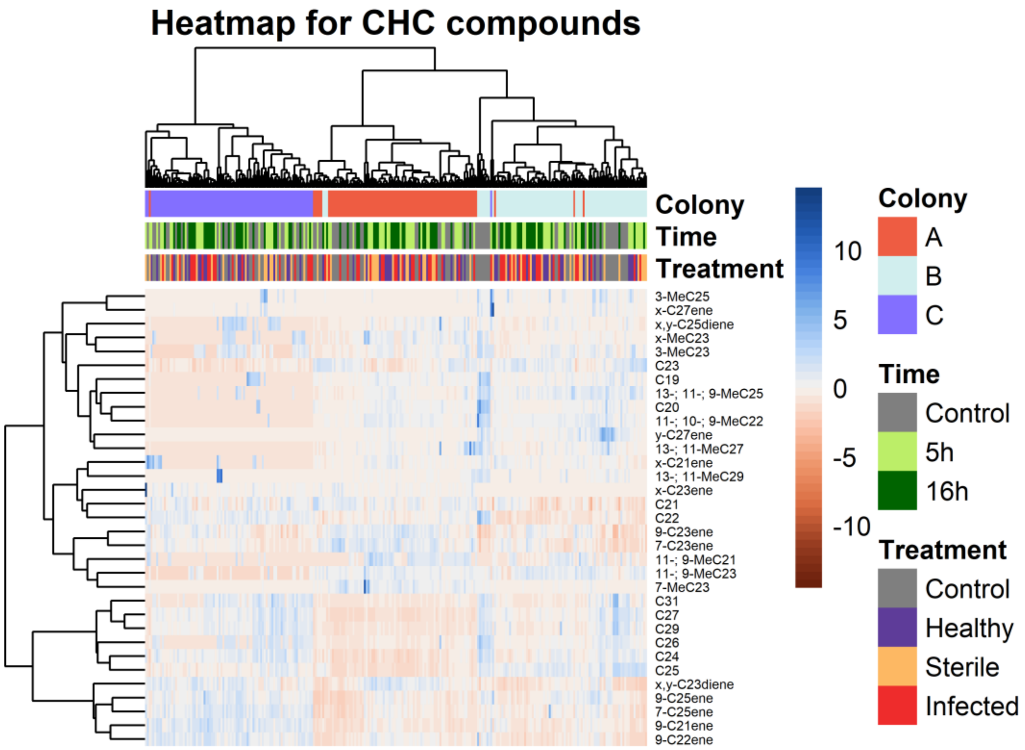


**Fig. S9. Heatmap visualising the intraspecific variation in CHC profiles**. Hierarchical clustering was performed on ants CHC profiles and their compounds (Z-scores) using the Ward's D2 method. Compounds whose structure could not be precisely identified are indicated with an x and y. Mean relative abundances and standard deviation among classes are shown in **Table S4**.

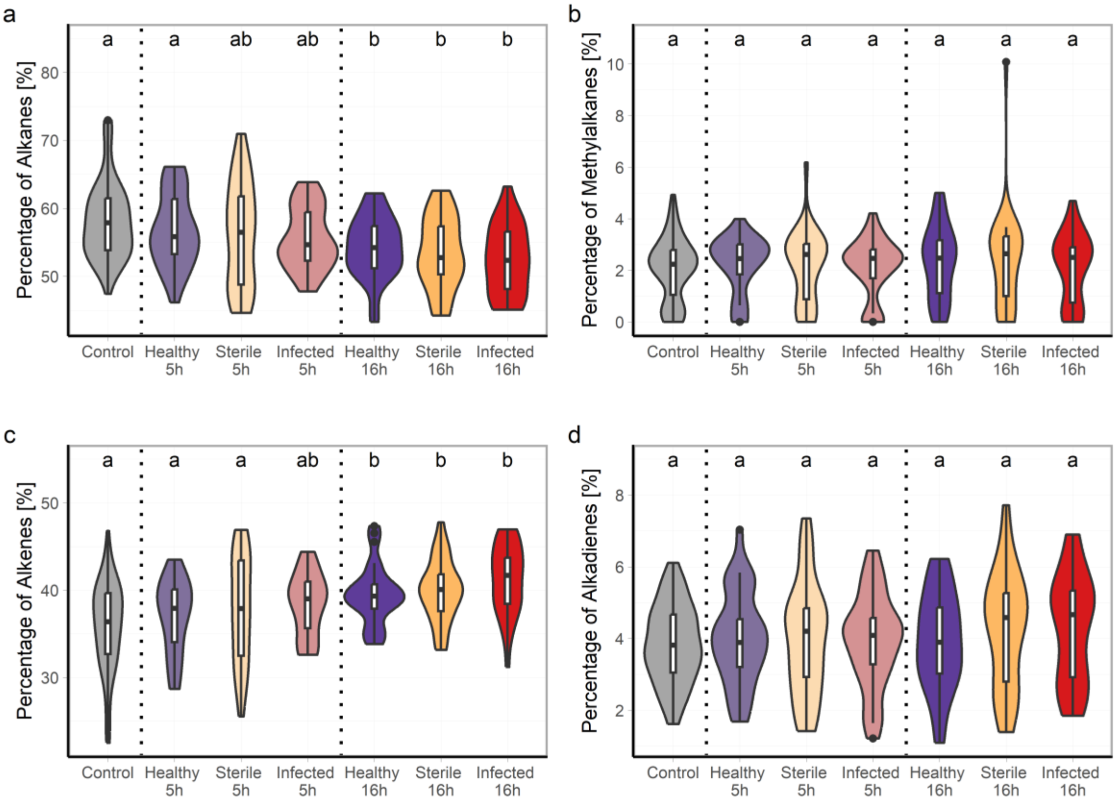


**Fig. S10. The percentage of each compound type in the CHC profile across treatments.** Compound types changed over time irrespective of injury condition (time after injury: PERMANOVA: F=15.52, df=2, $R^2$=0.11, p<0.001; treatment effect: F=1.09, df=2, $R^2$=0.01, p<0.001). The percentage of **(a)** Alkanes changed over time and injury conditions (GLM: *F* value (*F*)=7.67, df=6, p<0.001). The percentage of **(b)** Methylalkanes remained constant over time and injury condition (GLM: *F* value (*F*)=0.69, df=6, p=0.66), whereas the percentage of **(c)** Alkenes changed over time and injury condition (GLM: *F* value (*F*)=9.20, df=6, p<0.001). **(d)** Alkadienes showed no change in their relative abundance (GLM: *F* value (*F*)=0.977, df=6, p=0.44). Boxplots depict the first to third quartile of the interquartile range, horizontal lines within boxes are medians, and the whiskers are 1.5 interquartile range. Kernel densities are shown as shaded areas in violin plots. Generalised Linear Mixed Models were fitted for each compound type to examine the effects of time and injury condition. Colony ID was included as a random effect. Vertical dotted lines separate group treatments per timepoint. Different letters represent statistical significance for pair contrasts from estimated marginal means at p<0.05.

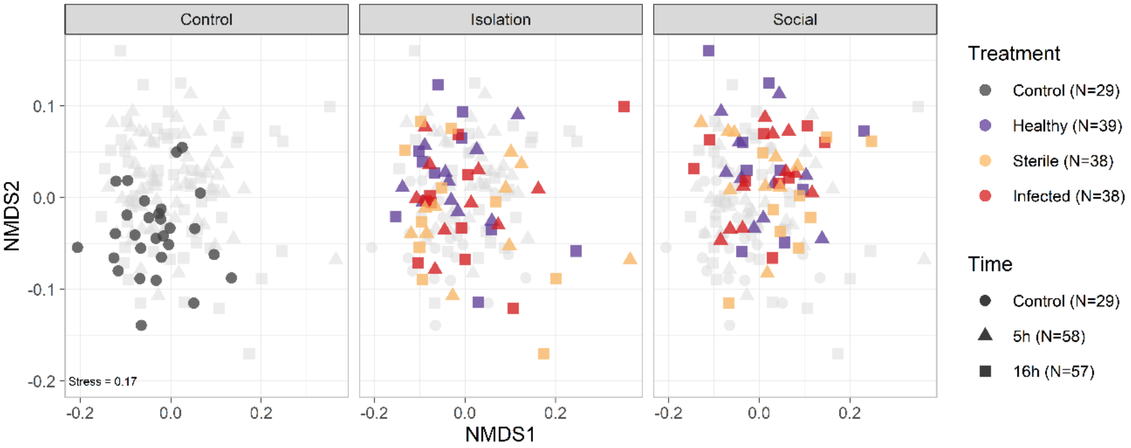


**Fig. S11. CHC profiles of ants that received infected or sterile wounds over time, either in isolation or with nestmates.** CHC profile dissimilarities are displayed as NMDS based on a Bray-Curtis dissimilarity matrix. Facets visualise data of ants after treatment (control), ants that remained in isolation (isolation), or ants that stayed with nestmates after treatment (social). Control (grey) corresponds to ants whose CHCs were extracted immediately after manipulation. Healthy (purple) were manipulated ants that did not receive an injury. Injured ants had their wounds exposed to either a pathogen solution (infected, red) or sterilised water (sterile, yellow). Shapes represent the time (hours) after manipulation. CHC profiles differed significantly between ants kept in a social or an isolated environment (**Table S7**). However, these differences were not associated with treatment type (**Table S8**). Survival after infection was similar to the other CHC experiment in isolation only (**Fig. S15, S16**).

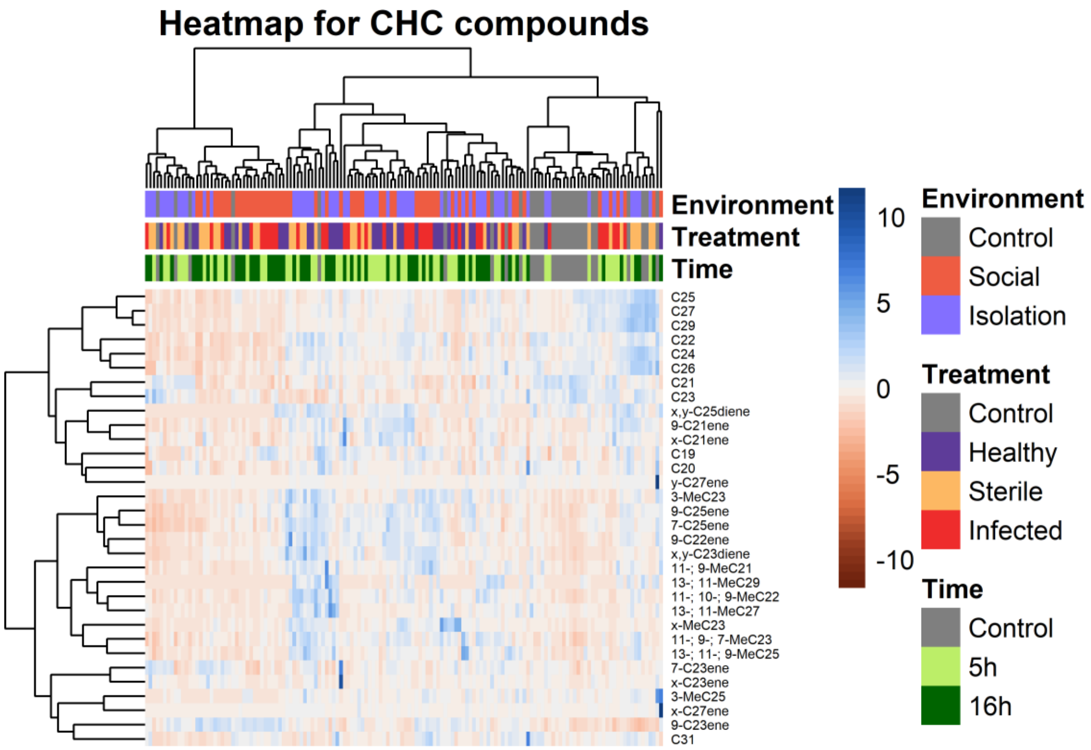


**Fig. S12. Heatmap visualising CHC compound differences across treatments, time and environment.** Hierarchical clustering was performed on ants CHC profiles and their compounds (Z-scores) using the Ward's D2 method. Compounds whose structure could not be precisely identified are indicated with an x and y. Mean relative abundances and standard deviation among classes are shown in **Table S9**.

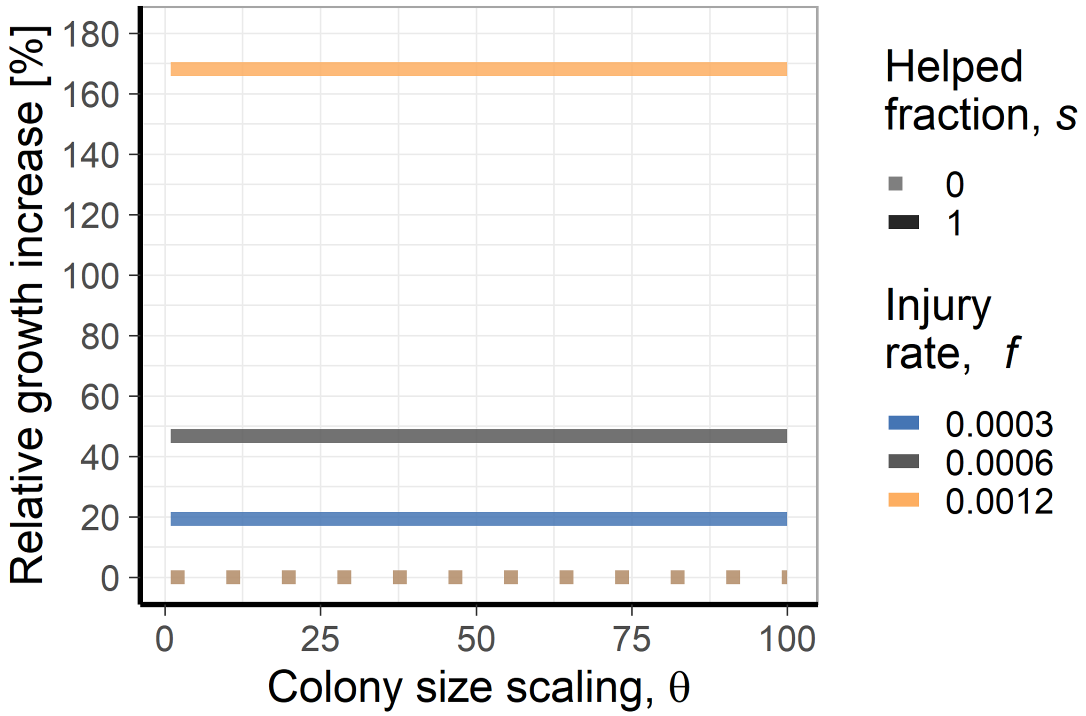


**Fig. S13. Effect of the colony size on colony growth under different injury ratios and degrees of care.** Relative colony growth increase is compared to colonies without care ($s$=0). The helping fraction $s$ represents the fraction of ants helped after each raid event in the discrete model. The Colony scaling $\theta$ factor represents species from a wide range of colony sizes (*e.g.* species with small colonies, such as *Megaponera analis* with 4000 ants, Colony scaling $\theta = 1$, to big colonies, such as *Eciton burchellii* with 400,000 ants, Colony scaling $\theta = 100$). The black line illustrates the growth rate dynamics for E. burchellii at the observed mean injury ratio (11%; $f$=0.0006). The effect of wound care on growth rates was greater when colonies had larger injury rates and were not affected at all by colony size.

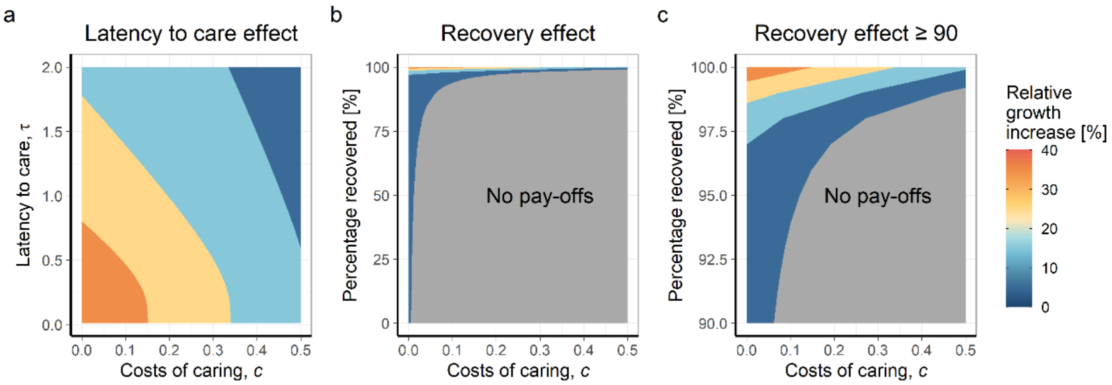


**Fig. S14. Effects on the colony growth rate when providing care incurs costs for the caregivers. (a)** The interaction between costs and the latency to care. As the costs of providing care increase, shorter latency is more important for maintaining high colony growth rates. Colours represent the change in the relative growth rate in bins of 10 units. Parameters used: Injury rate for *E. burchellii* $\boldsymbol{f}$=0.0006, $\boldsymbol{m_r}$=$\boldsymbol{m_i}$. **(b)** The interaction between the costs and the percentage of recovery after receiving care. A small increase in the cost of care strongly reduces colony growth and requires a high injury recovery rate after care to buffer these effects. Parameters used: $\boldsymbol{f}$=0.0006, $\boldsymbol{\tau}$=1. Colours represent the change in the relative growth rate in bins of 10 units. Grey area denotes the parameter space in which there is no benefit on colony growth rate (Growth rate $\boldsymbol{G \leq 0}$). **(c)** Detailed effect for recovery values greater than 90%.

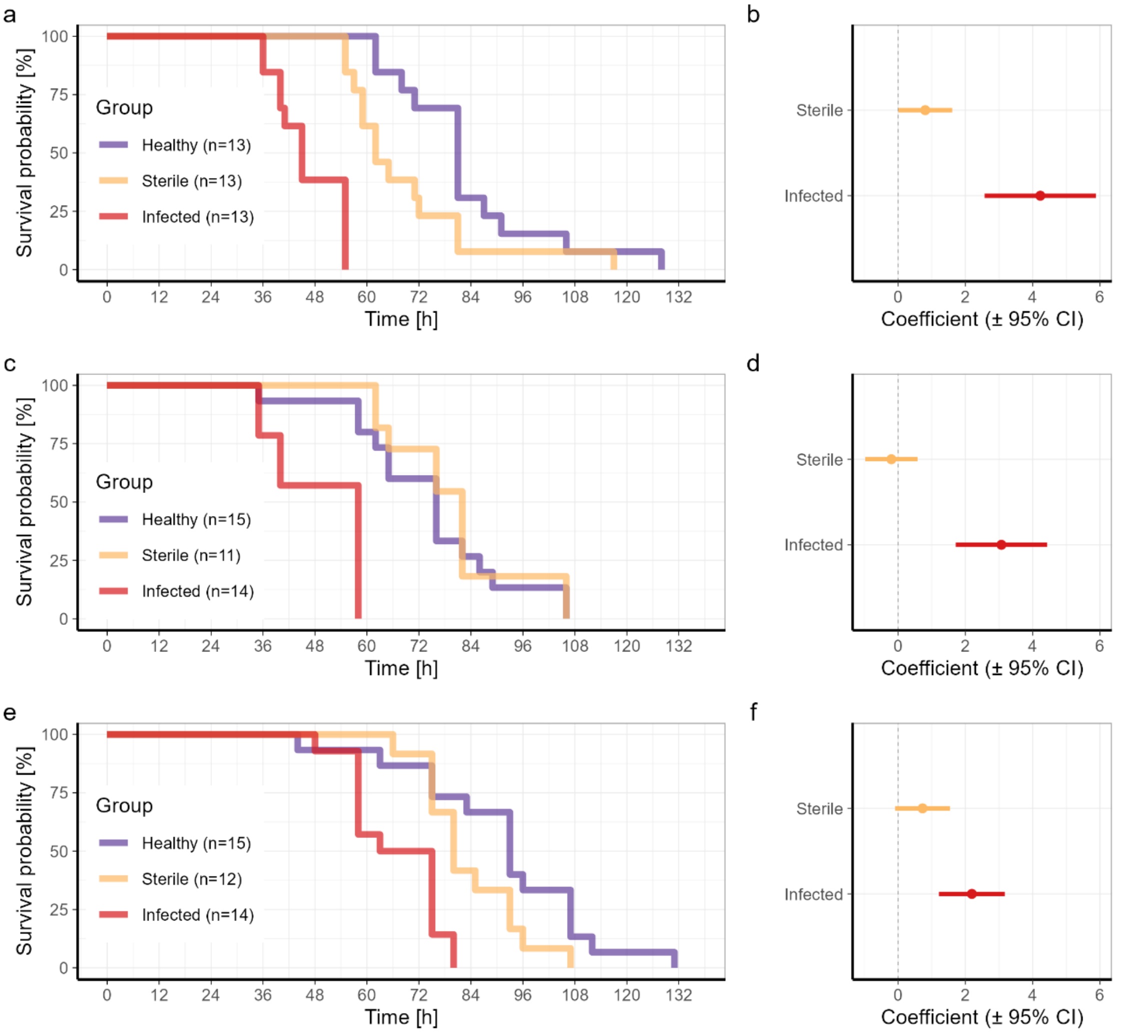


**Fig. S15. Survival curves for the three colonies used in the infection experiment for CHC analyses.** (**a & b**) Shows the Kaplan-Meier cumulative survival rates and Proportional Hazard ratio analysis for Colony A. (**c & d**) Shows the Kaplan-Meier cumulative survival rates and Proportional Hazard ratio analysis for Colony B. (**e & f**) Shows the Kaplan-Meier cumulative survival rates and Proportional Hazard ratio analysis for Colony C. In all cases, the Kaplan-Meier cumulative survival curves show that infected workers have lower survival than the healthy and sterile groups. Coefficient estimates of the Hazard ratios are on a logarithmic scale and the non-injured group (healthy) served as the reference (dashed vertical line). Proportional-hazard ratio analyses showed that infected workers exhibited significantly higher hazard ratios than healthy ants (**Table S10**).

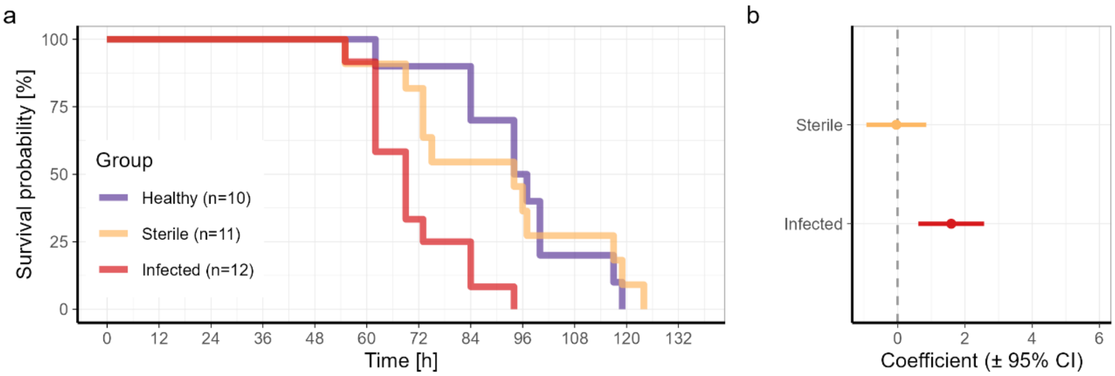


**Fig. S16. Survival curve for the colony used in the infection experiment for assessing the effect of the social environment on the change of CHCs. (a)** Shows the Kaplan-Meier cumulative survival rates. **(b)** Proportional Hazard ratio analysis. Coefficient estimates of the Hazard ratios are on a logarithmic scale. The non-injured group (healthy) served as the reference (dashed vertical line). Proportional-hazard ratio analyses showed that infected workers exhibited significantly higher hazard ratios than healthy ants (**Table S11**).

**Table S1. Pairwise comparisons for the injury ratio between body size classes.** *Post hoc* comparison conducted with a Dunn test.

| Comparison | z-statistic | p-value | p-adjusted |
|---|---|---|---|
| Media - Soldier | 5.838 | 0.000 | **<0.001** |
| Media - Minim | 2.289 | 0.011 | **0.017** |
| Soldier - Minim | -3.549 | 0.000 | **<0.001** |
| Media - Porter | 4.525 | 0.000 | **<0.001** |
| Soldier - Porter | -1.313 | 0.094 | 0.094 |
| Minim - Porter | 2.235 | 0.013 | **0.015** |

P-values below 0.05 are highlighted in bold after the Benjamini & Hochberg correction for multiple comparisons.

**Table S2. Contrast comparison for the ant survival from the experiment studying the effect of nest treatment after injury and infection in Fig. 2.** The survival of individuals was followed for 156 hours after isolation in sterile soil chambers. Survival functions were fitted with flexible parametric smooth generalised survival models. The model included fitted natural splines with three degrees of freedom to estimate the linear predictors of the survival function. The model included the colony as a stratum. No time-varying effects were assumed.

| Comparison | Estimate [CI: 95%] | z.ratio | p.value |
|---|---|---|---|
| Healthy - Infected+Wound care (12h) | 0.38 [0.24-0.6] | -4.17 | **<0.001** |
| Healthy - Infected | 0.14 [0.08-0.23] | -7.78 | **<0.001** |
| Healthy - Sterile+Wound care (12h) | 0.6 [0.39-0.93] | -2.29 | **0.032** |
| Healthy - Sterile | 0.65 [0.42-1.01] | -1.92 | 0.061 |
| Infected+Wound care (12h) - Infected | 0.37 [0.23-0.58] | -4.25 | **<0.001** |
| Infected+Wound care (12h) - Sterile+Wound care (12h) | 1.59 [1.01-2.51] | 2.01 | 0.055 |
| Infected+Wound care (12h) - Sterile | 1.71 [1.1-2.67] | 2.38 | **0.029** |
| Infected - Sterile+Wound care (12h) | 4.34 [2.67-7.08] | 5.89 | **<0.001** |
| Infected - Sterile | 4.68 [2.9-7.54] | 6.32 | **<0.001** |
| Sterile+Wound care (12h) - Sterile | 1.08 [0.69-1.68] | 0.32 | 0.745 |

P-values below 0.05 are highlighted in bold after the Benjamini & Hochberg correction for multiple comparisons

**Table S3. Hazard ratio estimates for ant survival in isolation from the experiment studying the effect of treatment at the bivouac after infections in Fig. S7.** Cox proportional hazard ratios were estimated for the effects of injury and infection over 162 hours for one colony. Coefficient estimates were exponentially transformed and represented with the 95% confidence interval (CI). Ants without any injury (healthy ants) were used as a reference group.

| Group | estimate [CI: 95%] | z.ratio | p.value |
|---|---|---|---|
| Sterile | 0.88 [0.47 - 1.64] | -0.41 | 0.683 |
| Infected | 11.91 [5.20 - 27.26] | 5.86 | **<0.001** |
| Infected + Treated | 0.81 [0.43 - 1.51] | -0.67 | 0.505 |

P-values below 0.05 are highlighted in bold.

**Table S4. Mean relative abundances (%) and standard deviation (sd) for all compounds observed in the CHC profile of ants according to the nest, treatment and time after treatment.** Compounds whose structure could not be precisely identified are indicated with an x and y.

| Compound | RI | Control | 5h | | | 16h | | |
|---|---|---|---|---|---|---|---|---|
| | | | Sterile | Infected | Healthy | Healthy | Sterile | Infected |
| C19 | 1,900 | 0.10 ± 0.11 | 0.09 ± 0.11 | 0.07 ± 0.07 | 0.05 ± 0.05 | 0.05 ± 0.04 | 0.06 ± 0.06 | 0.06 ± 0.06 |
| C20 | 2,000 | 0.06 ± 0.08 | 0.03 ± 0.03 | 0.04 ± 0.05 | 0.03 ± 0.03 | 0.03 ± 0.03 | 0.03 ± 0.03 | 0.03 ± 0.02 |
| 9-C21ene | 2,073 | 1.20 ± 0.63 | 1.34 ± 0.83 | 1.27 ± 0.70 | 1.28 ± 0.73 | 1.57 ± 0.72 | 1.36 ± 0.73 | 1.58 ± 0.82 |
| x-C21ene | 2,078 | 0.11 ± 0.15 | 0.10 ± 0.10 | 0.10 ± 0.15 | 0.06 ± 0.06 | 0.08 ± 0.08 | 0.06 ± 0.07 | 0.08 ± 0.09 |
| C21 | 2,100 | 20.92 ± 2.45 | 20.28 ± 2.66 | 20.41 ± 2.49 | 19.90 ± 2.02 | 19.53 ± 2.43 | 18.87 ± 3.02 | 19.37 ± 2.60 |
| 11-; 9-MeC21 | 2,138 | 0.09 ± 0.15 | 0.16 ± 0.16 | 0.18 ± 0.20 | 0.13 ± 0.19 | 0.13 ± 0.16 | 0.17 ± 0.18 | 0.07 ± 0.12 |
| 9-C22ene | 2,173 | 0.46 ± 0.17 | 0.46 ± 0.19 | 0.46 ± 0.17 | 0.46 ± 0.17 | 0.48 ± 0.17 | 0.49 ± 0.16 | 0.50 ± 0.17 |
| C22 | 2,200 | 2.42 ± 0.69 | 2.12 ± 0.54 | 2.08 ± 0.48 | 2.07 ± 0.44 | 1.95 ± 0.44 | 1.96 ± 0.41 | 1.87 ± 0.46 |
| 11-; 10-; 9-MeC22 | 2,234 | 0.09 ± 0.11 | 0.08 ± 0.07 | 0.07 ± 0.06 | 0.07 ± 0.05 | 0.08 ± 0.06 | 0.08 ± 0.06 | 0.07 ± 0.06 |
| x,y-C23diene | 2,267 | 3.75 ± 1.06 | 3.93 ± 1.29 | 3.90 ± 1.55 | 3.86 ± 1.26 | 3.85 ± 1.29 | 4.18 ± 1.56 | 4.22 ± 1.47 |
| 9-C23ene | 2,274 | 28.38 ± 4.52 | 29.43 ± 3.48 | 30.18 ± 4.83 | 30.48 ± 2.86 | 30.78 ± 2.87 | 31.33 ± 3.19 | 32.69 ± 3.30 |
| 7-C23ene | 2,280 | 2.71 ± 0.90 | 2.60 ± 0.59 | 2.73 ± 0.77 | 2.83 ± 0.46 | 2.98 ± 0.64 | 3.02 ± 0.69 | 3.23 ± 0.74 |
| x-C23ene | 2,291 | 0.18 ± 0.18 | 0.12 ± 0.13 | 0.30 ± 1.10 | 0.15 ± 0.30 | 0.13 ± 0.15 | 0.10 ± 0.13 | 0.10 ± 0.16 |
| C23 | 2,300 | 22.96 ± 2.63 | 22.72 ± 3.02 | 22.45 ± 4.21 | 22.54 ± 3.07 | 21.18 ± 3.01 | 21.65 ± 3.28 | 21.05 ± 3.18 |
| 11-; 9-MeC23 | 2,335 | 1.03 ± 0.66 | 1.24 ± 0.72 | 1.12 ± 0.78 | 1.18 ± 0.70 | 1.32 ± 0.88 | 1.25 ± 1.05 | 1.36 ± 0.98 |
| 7-MeC23 | 2,341 | 0.18 ± 0.22 | 0.15 ± 0.15 | 0.24 ± 0.42 | 0.17 ± 0.22 | 0.18 ± 0.30 | 0.26 ± 0.75 | 0.16 ± 0.25 |
| x-MeC23 | 2,363 | 0.13 ± 0.14 | 0.09 ± 0.07 | 0.11 ± 0.15 | 0.11 ± 0.12 | 0.12 ± 0.15 | 0.11 ± 0.14 | 0.06 ± 0.09 |
| 3-MeC23 | 2,372 | 0.24 ± 0.17 | 0.26 ± 0.17 | 0.22 ± 0.13 | 0.23 ± 0.13 | 0.24 ± 0.16 | 0.22 ± 0.15 | 0.20 ± 0.15 |
| C24 | 2,400 | 0.89 ± 0.28 | 0.86 ± 0.20 | 0.83 ± 0.29 | 0.82 ± 0.22 | 0.81 ± 0.20 | 0.79 ± 0.21 | 0.75 ± 0.22 |
| x,y-C25diene | 2,467 | 0.08 ± 0.12 | 0.08 ± 0.09 | 0.08 ± 0.10 | 0.08 ± 0.11 | 0.07 ± 0.12 | 0.09 ± 0.12 | 0.06 ± 0.10 |
| 9-C25ene | 2,473 | 1.76 ± 0.63 | 1.86 ± 0.34 | 1.64 ± 0.53 | 1.85 ± 0.54 | 2.16 ± 0.76 | 2.04 ± 0.67 | 1.85 ± 0.68 |
| 7-C25ene | 2,481 | 0.98 ± 0.32 | 1.01 ± 0.28 | 1.02 ± 0.51 | 1.04 ± 0.30 | 1.18 ± 0.34 | 1.16 ± 0.34 | 1.12 ± 0.34 |
| C25 | 2,500 | 7.28 ± 2.55 | 7.61 ± 2.60 | 7.22 ± 2.89 | 7.38 ± 2.10 | 7.55 ± 2.12 | 7.28 ± 2.47 | 6.84 ± 2.13 |
| 13-; 11-; 9-MeC25 | 2,531 | 0.14 ± 0.15 | 0.15 ± 0.13 | 0.11 ± 0.11 | 0.12 ± 0.09 | 0.15 ± 0.11 | 0.13 ± 0.12 | 0.11 ± 0.12 |
| 3-MeC25 | 2,573 | 0.08 ± 0.13 | 0.07 ± 0.11 | 0.09 ± 0.18 | 0.08 ± 0.23 | 0.07 ± 0.16 | 0.08 ± 0.20 | 0.02 ± 0.03 |
| C26 | 2,600 | 0.24 ± 0.19 | 0.16 ± 0.11 | 0.15 ± 0.13 | 0.15 ± 0.12 | 0.14 ± 0.11 | 0.12 ± 0.12 | 0.09 ± 0.09 |
| x-C27ene | 2,671 | 0.14 ± 0.31 | 0.09 ± 0.16 | 0.08 ± 0.19 | 0.23 ± 0.97 | 0.11 ± 0.28 | 0.38 ± 1.44 | 0.03 ± 0.07 |
| y-C27ene | 2,681 | 0.04 ± 0.06 | 0.02 ± 0.02 | 0.02 ± 0.05 | 0.02 ± 0.02 | 0.02 ± 0.04 | 0.01 ± 0.01 | 0.01 ± 0.02 |
| C27 | 2,700 | 1.74 ± 1.20 | 1.65 ± 0.89 | 1.65 ± 1.06 | 1.56 ± 0.94 | 1.78 ± 0.89 | 1.55 ± 0.96 | 1.41 ± 0.79 |
| 13-; 11-MeC27 | 2,731 | 0.06 ± 0.09 | 0.03 ± 0.03 | 0.03 ± 0.02 | 0.04 ± 0.07 | 0.04 ± 0.06 | 0.05 ± 0.11 | 0.03 ± 0.03 |
| C29 | 2,900 | 1.03 ± 0.86 | 0.95 ± 0.68 | 0.85 ± 0.70 | 0.75 ± 0.56 | 0.89 ± 0.55 | 0.80 ± 0.60 | 0.74 ± 0.60 |
| 13-; 11-MeC29 | 2,930 | 0.03 ± 0.07 | 0.02 ± 0.07 | 0.00 ± 0.01 | 0.00 ± 0.02 | 0.00 ± 0.01 | 0.00 ± 0.01 | 0.01 ± 0.02 |
| C31 | 3,100 | 0.52 ± 0.61 | 0.27 ± 0.35 | 0.29 ± 0.51 | 0.28 ± 0.37 | 0.36 ± 0.44 | 0.30 ± 0.37 | 0.23 ± 0.37 |

**Table S5. PERMANOVA analysis comparing CHC composition across injury type, time, and colony ID for the infection experiment.** The permutations (n = 9999) were not restricted to any group.

| Factor | Degrees of Freedom | Sum of Squares | $R^2$ | F | P-value |
|---|---|---|---|---|---|
| Group | 3 | 0.082 | 0.04 | 5.453 | **<0.001** |
| Time | 1 | 0.037 | 0.02 | 7.418 | **<0.001** |
| Colony | 2 | 0.865 | 0.38 | 86.220 | **<0.001** |
| Group:Time | 2 | 0.008 | 0.00 | 0.780 | 0.587 |

P-values below 0.05 are highlighted in bold.

**Table S6. Pairwise comparisons from PERMANOVA tests for the combination of injury condition and time after injury.** The false discovery rate was corrected by Benjamini & Hochberg's correction on the computed p-values (p-adjusted). The permutations (n = 9999) were restricted by Colony ID.

| Comparison | $R^2$ | p-value | p-adjusted |
|---|---|---|---|
| Control vs Healthy 16h | 0.042 | 0.0035 | **0.024** |
| Control vs Healthy 5h | 0.004 | 0.7291 | 0.837 |
| Control vs Infected 16h | 0.073 | 0.0001 | **0.002** |
| Control vs Infected 5h | 0.018 | 0.0913 | 0.274 |
| Control vs Sterile 16h | 0.045 | 0.0017 | **0.018** |
| Control vs Sterile 5h | 0.012 | 0.2080 | 0.352 |
| Healthy 16h vs Healthy 5h | 0.031 | 0.1291 | 0.305 |
| Healthy 16h vs Infected 16h | 0.026 | 0.1874 | 0.352 |
| Healthy 16h vs Infected 5h | 0.016 | 0.4437 | 0.666 |
| Healthy 16h vs Sterile 16h | 0.009 | 0.7570 | 0.837 |
| Healthy 16h vs Sterile 5h | 0.024 | 0.2180 | 0.352 |
| Healthy 5h vs Infected 16h | 0.068 | 0.0069 | **0.036** |
| Healthy 5h vs Infected 5h | 0.008 | 0.7558 | 0.837 |
| Healthy 5h vs Sterile 16h | 0.030 | 0.1305 | 0.305 |
| Healthy 5h vs Sterile 5h | 0.006 | 0.8315 | 0.862 |
| Infected 16h vs Infected 5h | 0.044 | 0.0420 | 0.176 |
| Infected 16h vs Sterile 16h | 0.009 | 0.7284 | 0.837 |
| Infected 16h vs Sterile 5h | 0.042 | 0.0557 | 0.195 |
| Infected 5h vs Sterile 16h | 0.014 | 0.4923 | 0.689 |
| Infected 5h vs Sterile 5h | 0.005 | 0.8623 | 0.862 |
| Sterile 16h vs Sterile 5h | 0.025 | 0.2067 | 0.352 |

P-values below 0.05 are highlighted in bold.

**Table S7. PERMANOVA analysis comparing CHC composition across environment (social or isolation), injury type and time.** Workers stayed in the artificial bivouac with other nestmates and larvae before extractions. The permutations (n = 9999) were not restricted to any group.

| Factor | Degrees of Freedom | Sum of Squares | $R^2$ | F | p-value |
|---|---|---|---|---|---|
| Environment | 2 | 0.170 | 0.24 | 22.826 | **<0.001** |
| Treatment | 2 | 0.007 | 0.01 | 0.995 | 0.416 |
| Time | 1 | 0.010 | 0.01 | 2.648 | **0.048** |
| Environment:Treatment | 2 | 0.004 | 0.01 | 0.554 | 0.773 |
| Environment:Time | 1 | 0.013 | 0.02 | 3.569 | **0.018** |
| Environment:Time | 2 | 0.001 | 0.00 | 0.113 | 0.997 |
| Environment:Treatment:Time | 2 | 0.006 | 0.01 | 0.847 | 0.525 |

P-values below 0.05 are highlighted in bold.

**Table S8. Pairwise comparisons from PERMANOVA tests for the combination of injury condition and time after injury for ants in a social environment.** Workers stayed in the artificial bivouac with other nestmates and larvae before extractions. The false discovery rate was corrected by Benjamini & Hochberg's correction on the computed p-values (p-adjusted). The permutations (n = 9999) were not restricted to any group.

| Comparison | R2 | p_value | p_adjusted |
|---|---:|---:|---:|
| Healthy 16h vs Healthy 5h | 0.111 | 0.0458 | 0.107 |
| Healthy 16h vs Infected 16h | 0.078 | 0.2498 | 0.375 |
| Healthy 16h vs Infected 5h | 0.119 | 0.1044 | 0.199 |
| Healthy 16h vs Sterile 16h | 0.055 | 0.3934 | 0.486 |
| Healthy 16h vs Sterile 5h | 0.217 | 0.0094 | **0.028** |
| Healthy 16h vs Control | 0.439 | 0.0001 | **0.000** |
| Healthy 5h vs Infected 16h | 0.079 | 0.1719 | 0.301 |
| Healthy 5h vs Infected 5h | 0.035 | 0.6642 | 0.697 |
| Healthy 5h vs Sterile 16h | 0.081 | 0.1874 | 0.303 |
| Healthy 5h vs Sterile 5h | 0.032 | 0.7569 | 0.757 |
| Healthy 5h vs Control | 0.326 | 0.0001 | **0.000** |
| Infected 16h vs Infected 5h | 0.062 | 0.3130 | 0.438 |
| Infected 16h vs Sterile 16h | 0.053 | 0.4524 | 0.512 |
| Infected 16h vs Sterile 5h | 0.154 | 0.0314 | 0.082 |
| Infected 16h vs Control | 0.361 | 0.0001 | **0.000** |
| Infected 5h vs Sterile 16h | 0.049 | 0.3838 | 0.486 |
| Infected 5h vs Sterile 5h | 0.044 | 0.4634 | 0.512 |
| Infected 5h vs Control | 0.243 | 0.0002 | **0.001** |
| Sterile 16h vs Sterile 5h | 0.133 | 0.0816 | 0.171 |
| Sterile 16h vs Control | 0.329 | 0.0001 | **0.000** |
| Sterile 5h vs Control | 0.264 | 0.0001 | **0.000** |

P-values below 0.05 are highlighted in bold.

**Table S9. Mean relative abundances (%) for all compounds observed in the CHC profile of ants according to the environment, treatment and time after treatment.** Workers in the "Social" group stayed in the artificial bivouac with other nestmates and larvae before extractions. Compounds whose structure could not be precisely identified are indicated with an x and y.

| Compounds | RI | Control | Isolation | | | | | | Social | | | | | |
|---|---|---|---|---|---|---|---|---|---|---|---|---|---|---|
| | | | 5h | | | 16h | | | 5h | | | 16h | | |
| | | | Healthy | Sterile | Infected | Healthy | Sterile | Infected | Healthy | Sterile | Infected | Healthy | Sterile | Infected |
| C19 | 1.900 | 0.04 | 0.03 | 0.04 | 0.04 | 0.04 | 0.03 | 0.04 | 0.05 | 0.05 | 0.05 | 0.04 | 0.04 | 0.06 |
| C20 | 2,000 | 0.04 | 0.03 | 0.03 | 0.04 | 0.03 | 0.04 | 0.03 | 0.02 | 0.03 | 0.03 | 0.03 | 0.02 | 0.03 |
| 9-C21ene | 2,081 | 1.02 | 1.33 | 1.35 | 1.76 | 1.31 | 1.56 | 1.93 | 1.31 | 1.45 | 1.26 | 0.88 | 1.02 | 0.91 |
| x-C21ene | 2,085 | 0.11 | 0.12 | 0.13 | 0.18 | 0.13 | 0.12 | 0.17 | 0.14 | 0.12 | 0.10 | 0.07 | 0.05 | 0.07 |
| C21 | 2,100 | 21.37 | 20.16 | 20.18 | 20.71 | 18.48 | 18.60 | 19.64 | 16.49 | 17.50 | 17.54 | 16.45 | 16.27 | 16.41 |
| 11-; 9-MeC21 | 2,137 | 0.11 | 0.15 | 0.11 | 0.13 | 0.13 | 0.17 | 0.14 | 0.13 | 0.13 | 0.16 | 0.14 | 0.09 | 0.21 |
| 9-C22ene | 2,173 | 0.22 | 0.24 | 0.21 | 0.27 | 0.23 | 0.27 | 0.25 | 0.28 | 0.32 | 0.24 | 0.22 | 0.20 | 0.25 |
| C22 | 2,200 | 0.97 | 0.93 | 0.76 | 0.90 | 0.87 | 0.88 | 0.79 | 0.81 | 0.81 | 0.78 | 0.70 | 0.66 | 0.79 |
| 11-; 10-; 9-MeC22 | 2,236 | 0.07 | 0.11 | 0.08 | 0.10 | 0.12 | 0.12 | 0.10 | 0.07 | 0.10 | 0.08 | 0.08 | 0.07 | 0.08 |
| x,y-C23diene | 2,267 | 0.53 | 0.62 | 0.50 | 0.66 | 0.69 | 0.82 | 0.59 | 0.86 | 0.96 | 0.77 | 0.77 | 0.60 | 0.86 |
| 9-C23ene | 2,274 | 38.92 | 40.97 | 40.57 | 40.46 | 40.68 | 39.60 | 40.09 | 44.80 | 43.25 | 44.36 | 48.23 | 46.47 | 46.23 |
| 7-C23ene | 2,280 | 3.19 | 3.74 | 3.58 | 3.61 | 3.47 | 3.72 | 3.75 | 4.02 | 3.99 | 3.48 | 3.33 | 3.48 | 2.76 |
| x-C23ene | 2,291 | 0.53 | 0.56 | 0.46 | 0.46 | 0.47 | 0.65 | 0.42 | 1.17 | 0.76 | 0.51 | 0.38 | 0.64 | 0.36 |
| C23 | 2,300 | 20.84 | 19.50 | 20.25 | 19.07 | 19.75 | 20.40 | 20.25 | 17.79 | 18.32 | 19.08 | 18.47 | 19.59 | 19.12 |
| 11-; 9-; 7-MeC23 | 2,335 | 1.06 | 1.62 | 1.31 | 1.46 | 1.82 | 1.80 | 1.58 | 1.29 | 1.49 | 1.29 | 1.25 | 1.16 | 1.27 |
| x-MeC23 | 2,358 | 0.36 | 0.75 | 0.33 | 0.48 | 0.68 | 0.49 | 0.38 | 0.43 | 0.54 | 0.49 | 0.31 | 0.31 | 0.59 |
| 3-MeC23 | 2,372 | 0.16 | 0.20 | 0.16 | 0.18 | 0.23 | 0.23 | 0.14 | 0.21 | 0.23 | 0.20 | 0.20 | 0.14 | 0.23 |
| C24 | 2,400 | 0.35 | 0.30 | 0.30 | 0.30 | 0.35 | 0.35 | 0.29 | 0.29 | 0.29 | 0.28 | 0.23 | 0.26 | 0.31 |
| x,y-C25diene | 2,467 | 0.01 | 0.01 | 0.01 | 0.01 | 0.01 | 0.01 | 0.01 | 0.01 | 0.01 | 0.00 | 0.01 | 0.00 | 0.01 |
| 9-C25ene | 2,474 | 1.05 | 1.15 | 1.05 | 1.23 | 1.44 | 1.23 | 1.07 | 1.65 | 1.65 | 1.36 | 1.41 | 1.16 | 1.54 |
| 7-C25ene | 2,481 | 0.72 | 0.80 | 0.63 | 0.85 | 0.94 | 0.83 | 0.76 | 1.03 | 1.01 | 0.77 | 0.78 | 0.68 | 0.92 |
| C25 | 2,500 | 6.46 | 5.29 | 6.17 | 5.65 | 6.28 | 5.96 | 5.90 | 5.91 | 5.54 | 5.81 | 4.91 | 5.76 | 5.74 |
| 13-; 11-; 9-MeC25 | 2,534 | 0.13 | 0.18 | 0.14 | 0.16 | 0.27 | 0.24 | 0.19 | 0.13 | 0.22 | 0.12 | 0.12 | 0.12 | 0.14 |

| Compounds | RI | Control | Isolation | | | | | | Social | | | | | |
|---|---|---|---|---|---|---|---|---|---|---|---|---|---|---|
| | | | 5h | | | 16h | | | 5h | | | 16h | | |
| | | | Healthy | Sterile | Infected | Healthy | Sterile | Infected | Healthy | Sterile | Infected | Healthy | Sterile | Infected |
| 3-MeC25 | 2,572 | 0.02 | 0.01 | 0.02 | 0.01 | 0.02 | 0.02 | 0.01 | 0.02 | 0.02 | 0.01 | 0.03 | 0.01 | 0.02 |
| C26 | 2,600 | 0.06 | 0.04 | 0.05 | 0.04 | 0.04 | 0.06 | 0.04 | 0.04 | 0.04 | 0.04 | 0.03 | 0.04 | 0.05 |
| x-C27ene | 2,667 | 0.04 | 0.03 | 0.02 | 0.03 | 0.04 | 0.03 | 0.03 | 0.03 | 0.04 | 0.02 | 0.16 | 0.02 | 0.03 |
| y-C27ene | 2,683 | 0.06 | 0.02 | 0.01 | 0.02 | 0.02 | 0.02 | 0.02 | 0.02 | 0.02 | 0.01 | 0.00 | 0.01 | 0.01 |
| C27 | 2,700 | 1.01 | 0.66 | 1.05 | 0.77 | 0.86 | 1.08 | 0.90 | 0.65 | 0.66 | 0.71 | 0.47 | 0.73 | 0.58 |
| 13-; 11-MeC27 | 2,731 | 0.17 | 0.20 | 0.13 | 0.16 | 0.34 | 0.20 | 0.14 | 0.08 | 0.18 | 0.11 | 0.08 | 0.11 | 0.16 |
| C29 | 2,900 | 0.26 | 0.16 | 0.29 | 0.19 | 0.17 | 0.29 | 0.24 | 0.16 | 0.16 | 0.20 | 0.11 | 0.20 | 0.11 |
| 13-; 11-MeC29 | 2,899 | 0.01 | 0.01 | 0.01 | 0.01 | 0.02 | 0.01 | 0.01 | 0.00 | 0.01 | 0.00 | 0.01 | 0.00 | 0.02 |
| C31 | 3,100 | 0.09 | 0.08 | 0.08 | 0.06 | 0.07 | 0.14 | 0.09 | 0.08 | 0.08 | 0.11 | 0.09 | 0.08 | 0.12 |

**Table S10. Hazard ratio estimates for ant survival from the experiment studying changes in the CHC profile after infection in Fig. S15.** Cox proportional-hazard ratios were estimated for the effects of injury and infection over 132 hours for the three colonies. Coefficient estimates were exponentially transformed and represented with the 95% confidence interval (CI). Ants without any injury (healthy ants) were used as a reference group. The likelihood ratio tests of models vs intercept-only models were: Colony A: $\chi2=40.11$, df=2, $p<0.001$; Colony B: $\chi2=31.67$, df=2, $p<0.001$; Colony C: $\chi2=20.4$, df=2, $p<0.001$.

| Group | estimate [CI: 95%] | z.ratio | p.value |
|---|---|---|---|
| *Colony A* | | | |
| Sterile | 2.25 [1.01 - 5.00] | 1.98 | **0.048** |
| Infected | 68.45 [13.05 - 358.91] | 5.00 | **<0.001** |
| *Colony B* | | | |
| Sterile | 0.82 [0.38 - 1.79] | -0.50 | 0.619 |
| Infected | 21.55 [5.54 - 83.78] | 4.43 | **<0.001** |
| *Colony C* | | | |
| Sterile | 2.07 [0.92 - 4.68] | 1.75 | 0.080 |
| Infected | 8.97 [3.37 - 23.84] | 4.39 | **<0.001** |

P-values below 0.05 are highlighted in bold.

**Table S11. Hazard ratio estimates for ant survival from the experiment studying changes in the CHC profile after infection in the social environment in Fig. S17.** Cox proportional-hazard ratios were estimated for the effects of injury and infection in the social environment over ~130 hours for one colony. Coefficient estimates were exponentially transformed and represented with the 95% confidence interval (CI). Ants without any injury (healthy ants) were used as a reference group. The likelihood ratio test of the model vs intercept-only model was: $\chi^2$=12.43, df=2, p<0.001.

| Group | estimate [CI: 95%] | z.ratio | p value |
|---|---|---|---|
| Sterile | 0.96 [0.40 - 2.34] | -0.09 | 0.932 |
| Infected | 4.92 [1.86 - 13.04] | 3.20 | **0.001** |

P-values below 0.05 are highlighted in bold.

**Movie S1 (separate file).** Antagonistic encounters between *Eciton burchellii* workers and *Odontomachus* sp., exemplifying potential injury events.

**Movie S2 (separate file).** Wound care (on-site care) is provided by nestmates in the field next to a raiding column.

**Movie S3 (separate file).** A nestmate collects secretions from its own metapleural gland and applies them to the wound of an infected worker inside an artificial nest.
